# Mating type specific transcriptomic response to sex inducing pheromone in the pennate diatom *Seminavis robusta*

**DOI:** 10.1101/2020.03.16.987719

**Authors:** Gust Bilcke, Koen Van den Berge, Sam De Decker, Eli Bonneure, Nicole Poulsen, Petra Bulankova, Cristina Maria Osuna-Cruz, Jack Dickenson, Koen Sabbe, Georg Pohnert, Klaas Vandepoele, Sven Mangelinckx, Lieven Clement, Lieven De Veylder, Wim Vyverman

**Affiliations:** Protistology and Aquatic Ecology, Department of Biology, Ghent University, 9000 Ghent, Belgium; Department of Plant Biotechnology and Bioinformatics, Ghent University, 9052 Ghent, Belgium; VIB Center for Plant Systems Biology, 9052 Ghent, Belgium; Department of Applied Mathematics, Computer Science and Statistics, Ghent University, 9000 Ghent, Belgium; Department of Statistics, University of California, Berkeley, Berkeley, CA, USA; SynBioC, Department of Green Chemistry and Technology, Ghent University, Coupure Links 653, 9000 Ghent, Belgium; B CUBE Center for Molecular Bioengineering, Technical University of Dresden, Tatzberg 41, 01307 Dresden, Germany; Bioinformatics Institute Ghent, Ghent University, Technologiepark 71, 9052 Ghent, Belgium; Marine Biological Association, The Laboratory, Citadel Hill, Plymouth PL1 2PB, UK; School of Ocean and Earth Science, University of Southampton, Southampton, United Kingdom; Bioorganic Analytics, Institute for Inorganic and Analytical Chemistry, Friedrich Schiller University Jena, Lessingstr. 8, D07743 Jena

**Author notes:** These authors contributed equally to this work (Gust Bilcke, Koen Van den Berge). Contact information of corresponding author Prof. dr. Wim Vyverman, Address: Krijgslaan 281 S8, 9000 Gent, Belgium.

## Abstract

Sexual reproduction is a fundamental phase in the life cycle of most diatoms. Despite its role as a source of genetic variation, it is rarely reported in nature and its molecular foundations remain largely unknown. Here, we integrate independent transcriptomic datasets, in order to prioritize genes responding to sex inducing pheromones (SIPs) in the pennate diatom *Seminavis robusta*. We observe marked gene expression changes associated with SIP treatment in both mating types, including an inhibition of S-phase progression, chloroplast division, mitosis and cell wall formation. Meanwhile, meiotic genes are upregulated in response to SIP, including a sexually induced diatom specific cyclin (dsCyc). Our data further suggest an important role for reactive oxygen species, energy metabolism and cGMP signaling during the early stages of sexual reproduction. In addition, we identify several genes with a mating type specific response to SIP, and link their expression pattern with physiological responses such as the production of the attraction pheromone diproline and mate-searching behaviour in MT+. Combined, our results provide a model for early sexual reproduction in pennate diatoms and significantly expand the suite of target genes to detect sexual reproduction events in natural diatom populations.

## Introduction

Sexual reproduction is a virtually universal feature in the life cycle of eukaryotic organisms and a wealth of reproductive strategies has evolved across different phyla^1^. Likewise, sexual reproduction is an essential phase in the diplontic life cycle of most diatoms, an extraordinarily diverse group of microalgae that play an important role in primary production and biogeochemical cycling in the oceans^2,3^. Their unique life cycle is characterized by cell size reduction during vegetative growth^4^. Cells become sexually active once their size is below a species-specific sexual size threshold (SST). Sexual reproduction restores the maximum cell size by expansion of the zygote to form large auxospores that release an initial cell^4^. Although the conservation of meiotic genes^5^ and population genetic data on sexual homologous recombination ^6,7^ suggest that sexual reproduction occurs in natural diatom populations, reports on sexual events remain scarce and are usually restricted to field sites with high frequency monitoring^8–11^. Successful crossing of diatoms in laboratory conditions, however, revealed a diverse range of life cycle strategies with unique features for centric, araphid pennate and raphid pennate diatoms^4,12^.

Characteristic for pennate diatoms, sexual reproduction is initiated by the interaction of sexually mature vegetative cells (gametangia) from compatible mating types^4,13^. Whereas for planktonic species mainly passive physical forces are thought to steer cell-cell interaction^14^, most benthic raphid diatoms actively move towards a partner of the compatible mating type^13^. Recently, experimental evidence has shown that certain pennate diatoms deploy sex pheromones to recognize and localize a suitable partner^15–18^. Furthermore, a multistage pheromone cascade was discovered in the raphid pennate diatom *Seminavis robusta* (Figure 1), emphasizing the largely unexplored complexity of mate localization and recognition in diatoms^17^. Previous studies have briefly addressed the transcriptomic response to sex inducing pheromones (SIPs) in pennate diatoms^17,18^, but a detailed overview and timing of expression is currently lacking. Importantly, while the previously discovered multistage pheromone cascade suggests a mating type specific response to SIPs, it is unknown how this is reflected at the molecular level.

**Figure 1:**
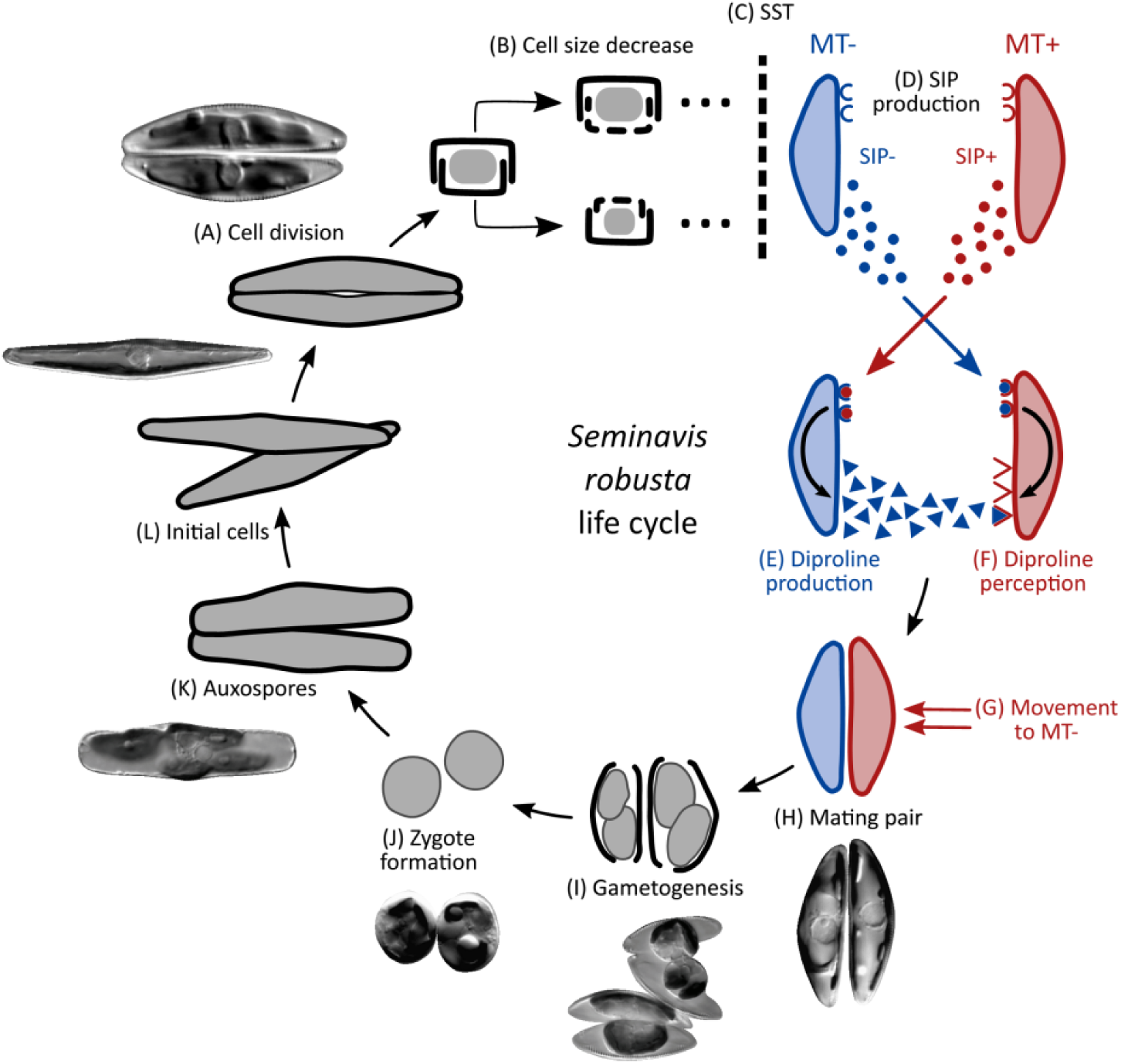
The life cycle of *Seminavis robusta.* The *S. robusta* life cycle is diplontic and consists of long periods of vegetative division alternating with short periods of sexual reproduction. **(A)** The average cell size decreases with every mitotic division. **(B)** Transverse view of a vegetative cell showing the mechanism of cell size decrease. **(C)** When populations pass the sexual size threshold (SST) at a cell size of 50 µm, cells become capable of sexual reproduction. **(D)** Mating type + (MT+) and mating type - (MT-) start producing sex inducing pheromones called SIP+ and SIP-, respectively. SIP induces a cell cycle arrest in the compatible mating type. **(E)** In response to SIP+, MT- secretes an attraction pheromone: the diketopiperazine diproline, while **(F)** MT+ becomes sensitive to diproline and glides towards the diproline source. **(G, H)** Diproline signaling leads to mate finding and pair formation. **(I, J)** Each partner produces two gametes that fuse with the gametes of the compatible mating type to form zygotes. Finally, auxospores **(K)** will enlarge and release an initial cell of the original cell size **(L)**. The molecular basis underlying the life cycle of pennate diatoms is to date largely unexplored. This figure was modified from Moeys *et al.* (2016) and Gillard *et al.* (2012) and some microscopic pictures were obtained from Chepurnov *et al.* (2002).

Here, we use RNA-sequencing to provide the first detailed description of the response to SIPs in *S. robusta.* Responses in gene expression were compared between existing datasets for mating type - (MT-) and a newly generated time-series RNA-seq dataset for the compatible mating type MT+. We relate physiological changes resulting from a G1 phase arrest to differentially expressed genes throughout the cell cycle and show that SIP- increases motility of the attracted MT+. To tackle technical challenges in comparing gene expression among mating types, we introduce a workflow to integrate RNA-seq datasets which allows a comparison between independent datasets. This approach allowed the identification of key genes exhibiting either mating type specific or shared responses to SIP. These key genes include a sexually induced cyclin and highlight the importance of reactive oxygen species (ROS), energy metabolism and ubiquitination in the mating process.

## Material and methods

### Culture conditions

*Seminavis robusta* strains were obtained from the Belgian Coordinated Collection of Microorganisms (BCCM/DCG) and were grown in sterile filtered natural sea water (NSW) from the North Sea enriched with Guillard’s F/2 solution, except for the experiments involving MT+ motility and MT- flow cytometry where cultures were grown in artificial sea water (ASW) with F/2 solution. Prior to the experiments, cultures were made axenic by adding 500 mg/L penicillin, 500 mg/L ampicillin, 100 mg/L streptomycin and 50 mg/L gentamycin to the medium. Cultures were grown at 18 °C in 12h:12h light:dark cycles under cool white fluorescent lamps unless stated otherwise.

### Preparation of SIP containing filtrate

MT- cultures (strain 85B) and MT+ cultures (strain D6) below the SST were cultured in 150mL cell culture flasks for one week. When cultures reached the late-exponential growth phase the medium was vacuum filtered using a Stericup with pore size of 0.22 µm (Merck GmbH, Germany) in order to obtain a filtrate containing SIP- and SIP+, respectively. The potency of the filtrate was assessed using a cytokinetic arrest and a diproline attraction assay (Supplementary Methods). The SIP filtrate was used for the RNA-seq experiment, for cell cycle analysis using flow cytometry and for assessing its effect on motility of MT+ (Supplementary Methods).

### RNA-seq experimental setup, data analysis and functional interpretation

RNA-seq data generated in this study representing the response of MT+ to SIP- was complemented with existing data on the response of MT- to SIP+. The experimental setup for the two MT- RNA-seq datasets are described in their respective publications^17,19^. For the novel MT+ dataset, we harvested control cultures in the dark (time point 0h) and further harvested dark-synchronized cultures consisting of three control replicates and three SIP- treated replicates at five time points: 15min, 1h, 3h, 6h and 9h. Details about the experimental setup are described in Supplementary Methods.

All three RNA-seq datasets were mapped with Salmon v0.9.1^20^ to the *S. robusta* gene models v1.0 from Cirri et al.^19^ (available at https://bioinformatics.psb.ugent.be/gdb/seminavis/Version1.0; mapping rates shown in Supplementary Figure 1) and a conventional differential expression (DE) analysis was performed using negative binomial models implemented in the edgeR package^21^. Details about the model, design matrix and contrasts of interest are described in Supplementary Methods. Functional annotation for all genes was derived using an ensemble of three methods: InterProScan^22^, AnnoMine^23^ and eggNOG-mapper^24^. Gene families were computed by clustering protein sequences with TRIBE-MCL^25^. Specific details on the functional annotation generation are described in Supplementary Methods.

To identify biologically relevant DE genes, we used a two-fold approach. First, we used the results from the conventional DE analysis to look into genes and biological processes involved in sexual reproduction, based on the genes’ functional annotation and the current literature. Second, we developed a novel integrative workflow that allowed us to compare the response to SIP in different RNA-seq datasets from both mating types. We restricted the comparison to the time points that are available for both mating types (15min, 1h and 3h). Genes discovered by the integrative analysis represent key genes with a shared versus mating type specific response to SIP. Three sets of genes were defined: genes responding to SIP in both mating types (SRBs: “SIP Responsive in Both mating types”), genes with a MT- specific response (SRMs: “SIP Responsive in mating type Minus”) and genes with a MT+ specific response (SRPs, “SIP Responsive in mating type Plus”). Details about the integrative workflow can be found in Supplementary Methods.

## Data availability and reproducibility

The raw data from the new RNA-seq experiments are available at the European Nucleotide Archive (ENA) at EMBL-EBI under accession number PRJEB35793 (https://www.ebi.ac.uk/ena/data/view/PRJEB35793). All scripts required to reproduce the analyses and figures reported in this paper as well as Salmon estimated count matrices and DE analysis results are available on our GitHub repository at https://github.com/statOmics/SeminavisComparative.

## Results and discussion

In this study, we generated a new time-series dataset to investigate the response of dark-synchronized MT+ cultures to SIP- (6 time points, 0-9h, Figure 2A) and to compare expression patterns with two existing datasets on the response of MT- to SIP+ (5 time points, 0-10h, Figure 2B)^17,19^. For each dataset, three replicates are available for both control and SIP treated conditions at every time point. Here, we first report the results of a separate differential expression (DE) analysis for each dataset. Next, we use these results to discover genes or biological processes related to sexual reproduction, either based on their predicted functional annotation or the current literature. Finally, we integrate the results of both the novel and publicly available datasets. However, directly comparing sets of differentially expressed genes between datasets is inappropriate. Indeed, due to the inherent low power to detect true DE genes in RNA-seq experiments, a substantial proportion of false negative genes is expected among the non-significant gene list. This invalidates an approach that would depict genes that are only significantly DE in one mating type as uniquely responding in that mating type. We tackled this problem by developing a statistical integrative analysis framework that is capable of testing for equivalent, i.e. non-DE, expression between conditions. Coupling equivalence testing in one MT with DE calls in the other allowed for the discovery of key genes exhibiting mating type specific responses to SIP, as well as key genes responsive in both mating types.

**Figure 2:**
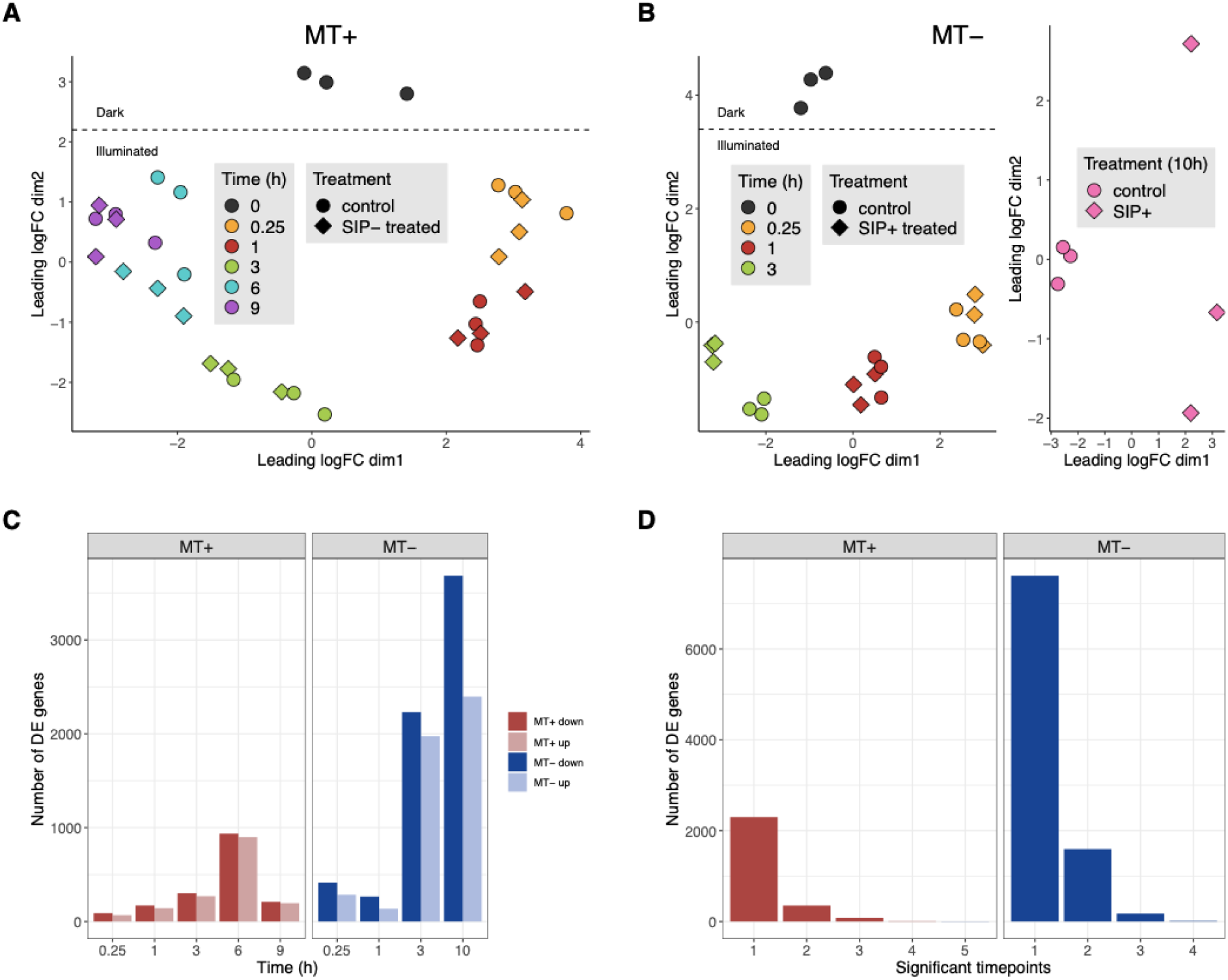
Transcriptional responses induced by SIP treatment for both mating types of *Seminavis robusta*. (**A**) Multidimensional scaling (MDS) plot of time series for MT+ expression data (0h-9h), and (**B**) MDS plots of two MT- expression datasets (0h-3h and 10h, respectively). Distances between samples in the MDS plot approximate the log2 fold change of the top 500 genes. (**C**) Number of DE genes between control and SIP treated cultures for each time point in both mating types. Each dataset was analyzed on a 5% overall FDR (OFDR) level, i.e. the fraction of false positive genes over all rejected genes. The colour of bars represents mating type (MT+: red, MT-: blue) and the direction of change (dark: downregulated after SIP treatment, light: upregulated after SIP treatment). (**D**) The number of time points each DE gene is significant at. The colour of bars represents mating type (MT+: red, MT-: blue). Most genes are significant in only a few time points, whereas a smaller number of genes are significantly differentially expressed in multiple time points.

### General transcriptional response and identification of key SIP responsive genes

Multidimensional scaling plots of the RNA-seq data (Figure 2A-B) showed that in both mating types the dark-to-light transition and time since illumination were the major drivers of gene expression change throughout the experiments. However, as time progresses, the effect of SIP becomes more pronounced. This is supported by differential expression analysis showing that the number of significant genes increased markedly at later time points (Figure 2C). Overall, most genes are significant in only a single time point (5% OFDR, Figure 2D). Combined, on a 5% overall FDR (OFDR) level, 4037 genes are DE in response to SIP treatment for MT+, while 5486 genes are found to be DE in MT- in the first three hours ^17^ and 6079 genes after 10h^19^.

Gene sets involved in amongst others cell cycle, meiosis and cyclic nucleotide signalling were enriched in both mating types, while proline biosynthetic process gene set was enriched uniquely in MT-(Supplementary Figure 2), in accordance with findings in previous studies^17,18^. Figure 3 plots genes involved in these processes that were significantly DE in the conventional DE analysis.

**Figure 3:**
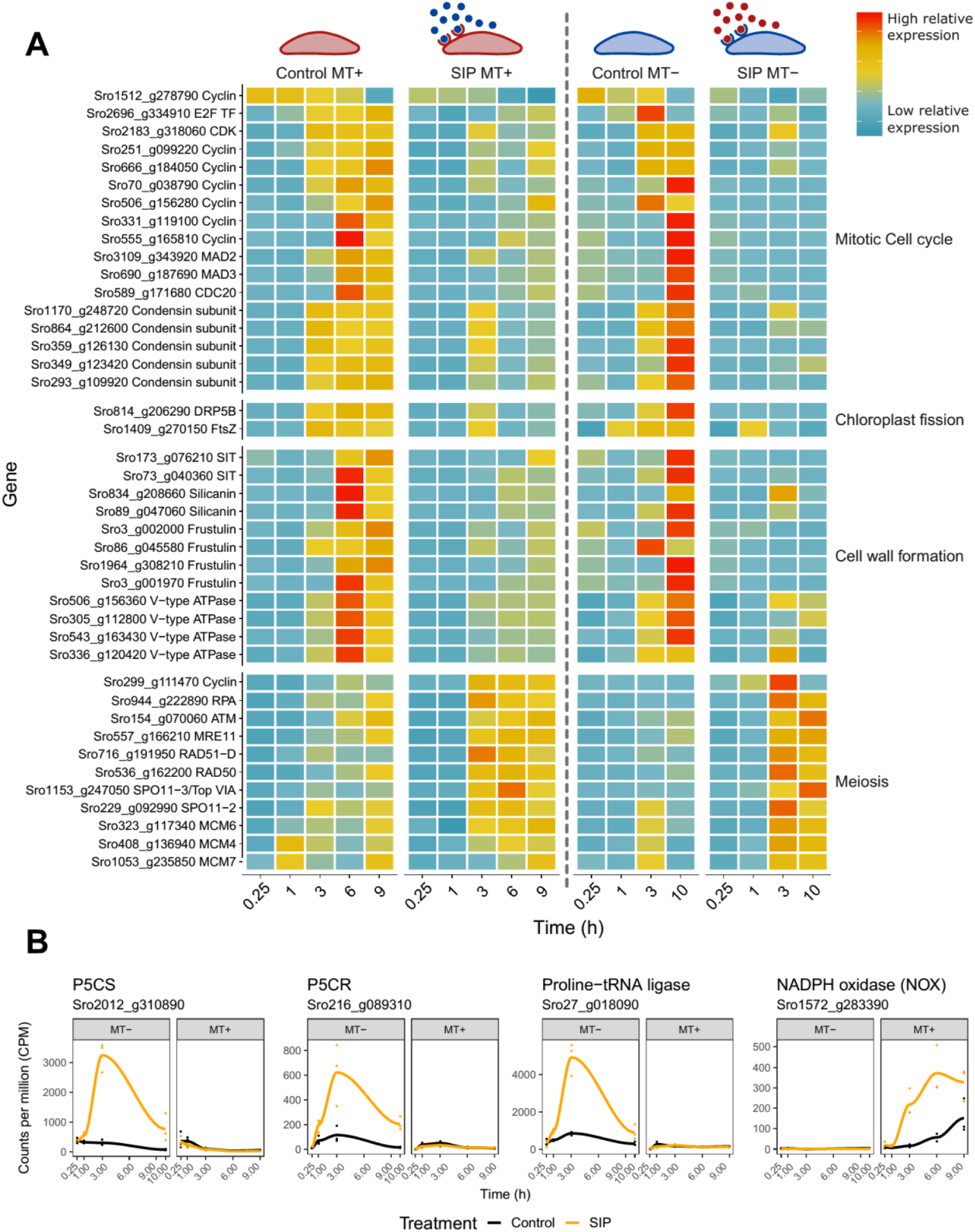
Gene expression of *Seminavis robusta* genes involved in mating related processes. **(A)** Heatmap of genes related to mitotic and meiotic cell cycle progression which are upregulated in both mating types in the conventional differential expression (DE) analysis. Each gene is plotted for control and SIP treated conditions in both *S. robusta* mating types. Genes are specified as row names and are coded according to their expression, amounting to counts per million (CPM) standardized to zero mean and unit variance for each gene in each mating type separately. Blue indicates low expression, while red indicates high expression. **(B)** Expression of genes related to diproline synthesis and reactive oxygen species (ROS) production which are significantly DE in only one mating type in the conventional DE analysis. CPM are plotted as a function of time for both mating types. The data points correspond to gene expression of the replicates in each time point while the solid line represents the mean. Data points and lines are colored according to condition, i.e. black for control condition and orange for SIP treatment. P5CS = Δ1-pyrroline-5-carboxylate synthetase; P5CR = Δ1-pyrroline-5-carboxylate reductase.

Complementary, our integrative workflow revealed 52 key genes responding to SIP in both mating types (SRBs), 12 genes uniquely responding in MT+ (SRPs) and 70 genes uniquely responding in MT- (SRMs) (Figure 4A, Supplementary Figures 3-5). Remarkably, while in MT- we discovered a comparable number of down- and upregulated SRMs, we only find upregulated SRPs and SRBs, indicating that sexual processes induced by SIPs are mainly driven by the induction of key genes rather than the downregulation of inhibitory genes (Figure 4B).

**Figure 4:**
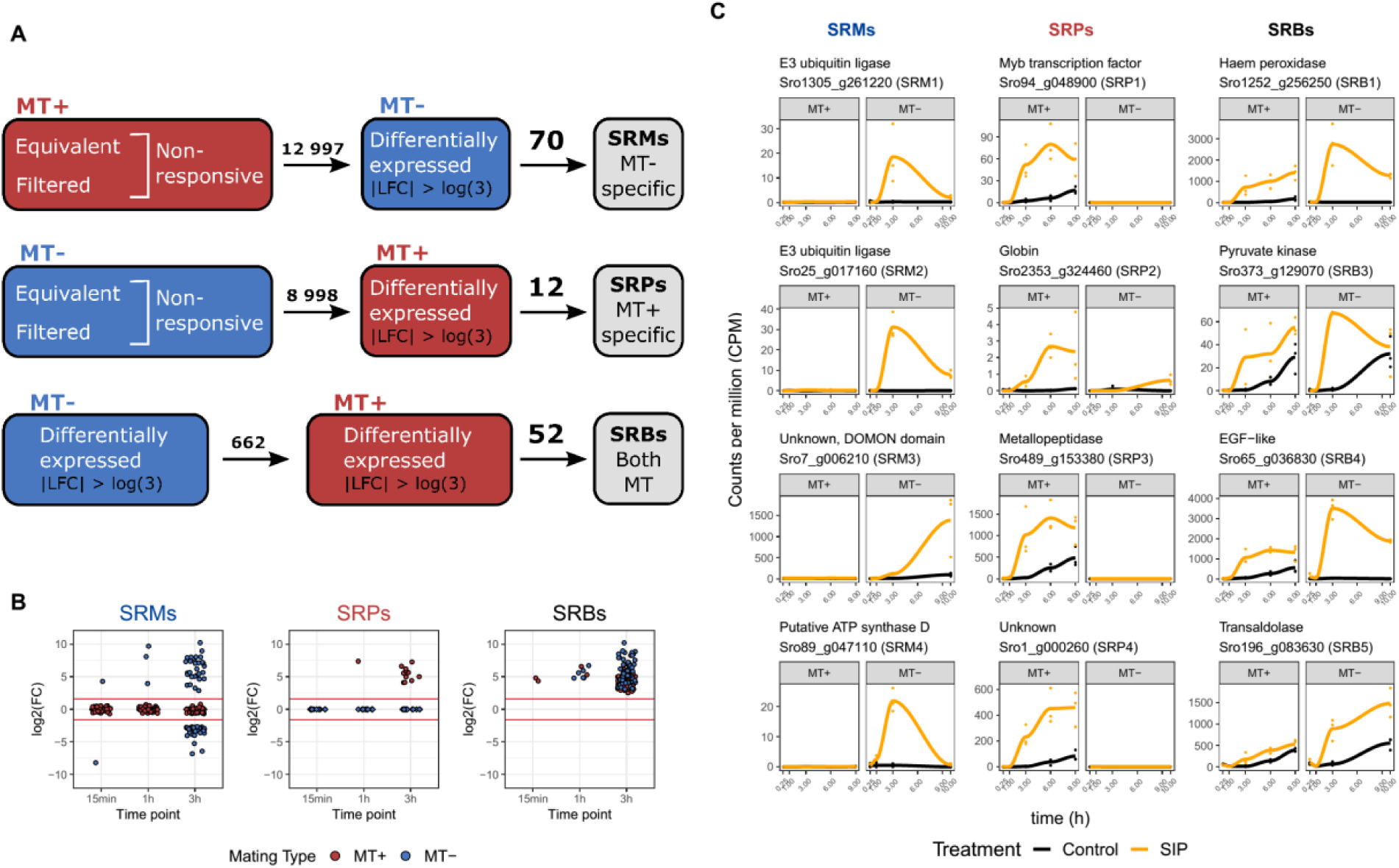
Visualization and main results of the integrative workflow. **(A)** Schematic representation of the integrative data analysis indicating how SIP responsive genes with a shared response (SRBs) or mating type specific response (SRPs, SRMs) were discovered. Numbers above the arrows indicate the number of genes withheld after each step. Non-responsive genes in one mating type were defined as the union of equivalent and filtered (lowly expressed) genes. A log fold change (LFC) cutoff of +- log(3) was used to define differentially expressed genes and equivalent genes. **(B)** Log2 fold changes of SRMs, SRPs and SRBs in both mating types. Each gene is plotted for the time point at which they are differentially expressed. Genes which were not expressed (“filtered”) in the non-responsive mating type are plotted as diamonds. The red horizontal lines represent the fold change cutoff used to determine equivalence and differential expression. **(C)** Expression of a selection of SIP responsive genes (SRMs, SRPs, SRBs). For each gene, counts per million (CPM) are plotted as a function of time for both mating types. The data points correspond to gene expression of the replicates in each time point and the solid line represents the mean. Data points and lines are colored according to condition, i.e. black for control condition and orange for SIP treatment.

In what follows we will discuss in more detail the genes and pathways that are responding to SIP in both mating types or uniquely in one mating type and link these changes to physiological events in the mating process.

### Responses to SIP conserved in both mating types

#### Integrative analysis reveals key genes responsive in both mating types

A large fraction (22/52) of SRBs - key genes with a strong response to SIP in both mating types - lack any functional annotation and homology to sequenced genomes of other diatoms (Supplementary Table 1), suggesting that the molecular mechanisms underlying early mating are highly species-specific. The remaining 30 SRBs can be linked to energy metabolism, ROS signaling and meiosis. Pyruvate kinase (*Sro373_g129070*) and isocitrate dehydrogenase (*Sro492_g153950*), respectively involved in glycolysis and the citric acid cycle, are strongly upregulated in both mating types (Figure 4C), suggesting an increased energy demand. Interestingly, pyruvate kinase is also upregulated during gametogenesis in the brown alga *Saccharina latissima*^*26*^ and the parasite *Plasmodium berghei*^*27*^. Additionally, two enzymes from the pentose phosphate pathway (PPP) are among the SRBs: transketolase (*Sro524_g159900*) and transaldolase (*Sro196_g083630*) (Figure 4C). The PPP generates NADPH, a reductive compound needed in various metabolic reactions and involved in detoxification of ROS by regenerating glutathione^28,29^. Furthermore, one SRB encoding a haem peroxidase (*Sro1252_g256250*) exhibited strong upregulation upon SIP treatment (Figure 4C). Upregulation of haem peroxidases was also reported during sexual reproduction in other eukaryotes, e.g. mosquitoes (*Anopheles gambiae*)^30^ *and fungi*^*31,32*^. *Haem peroxidases promote substrate oxidation in various metabolic pathways and are essential for the detoxification of ROS*^*33*^, *suggesting that ROS signaling plays a role in the response to SIP, as seen in the green algae Volvox carteri*, where high ROS levels trigger sex^34^. Finally, a highly expressed SRB encodes a transmembrane protein containing an Epidermal Growth Factor-like (EGF-like) domain (*Sro65_g036830*, Figure 4C). EGF-like domains are generally extracellular protein modules that play a role in receptor/ligand interactions, intracellular signaling and adhesion^35^. Since membrane bound proteins containing EGF-like domains are required for gamete fusion in green algae^36^ and oocyte binding in animals^37–39^, we speculate that in *S. robusta*, this gene plays a role in cell-cell communication between gametangia during mate pairing or in the recognition and fusion of gametes.

#### Conventional DE analysis uncovers mating-related processes differentially expressed in both mating types

We used flow cytometry to confirm a sex pheromone induced G1 phase arrest, which was proposed for *S. robusta* and *Pseudo-nitzschia multistriata*^*17,18*^. Treatment with SIP significantly decreased the proportion of G2/M phase cells in MT+ after 3h (control 12.4% versus SIP 3.5%, p = 0.0023) and 9h (control 20.9% versus SIP 9.4%, p = 0.0123) and after 9h in MT- (control 10.5% versus SIP 3.3%, p < 0.0001) (Figure 5C). A temporary arrest of the cell cycle appears to be a prerequisite for a switch to meiosis in many diatoms^17,18,40,41^, although in the centric diatom *Skeletonema marinoi* no growth arrest was observed during sexual reproduction^42^. The cell cycle arrest is reflected in the transcriptomic data as a sequential downregulation of cell cycle genes in the general DE results (Figure 3). In eukaryotes, S-phase progression is controlled by E2F transcription factors forming a heterodimer with Dimerization Partner (DP) transcription factors^43^. In accordance with plants and animals^44,45^, we observe an increase in expression of E2F transcription factors in control conditions as cells go through the cell cycle. SIP treatment significantly repressed expression of two E2Fs (*Sro2696_g334910* and *Sro1798_g298290*) in both mating types and MT-, respectively (Supplementary Figure 6). The transcriptional repression of E2Fs is likely caused by the inability of cells to enter S-phase as a result of the G1 phase arrest, although the activity of E2Fs is generally also regulated by other mechanisms such as Rb-related protein binding and phosphorylation which we did not assess^44,45^. Furthermore, two DP genes were DE in MT- in response to SIP+; one (*Sro905_g218540*) was repressed, while the other (*Sro905_g218570*) was induced, suggesting they play a contrasting role in the cell cycle arrest (Supplementary Figure 6).

**Figure 5:**
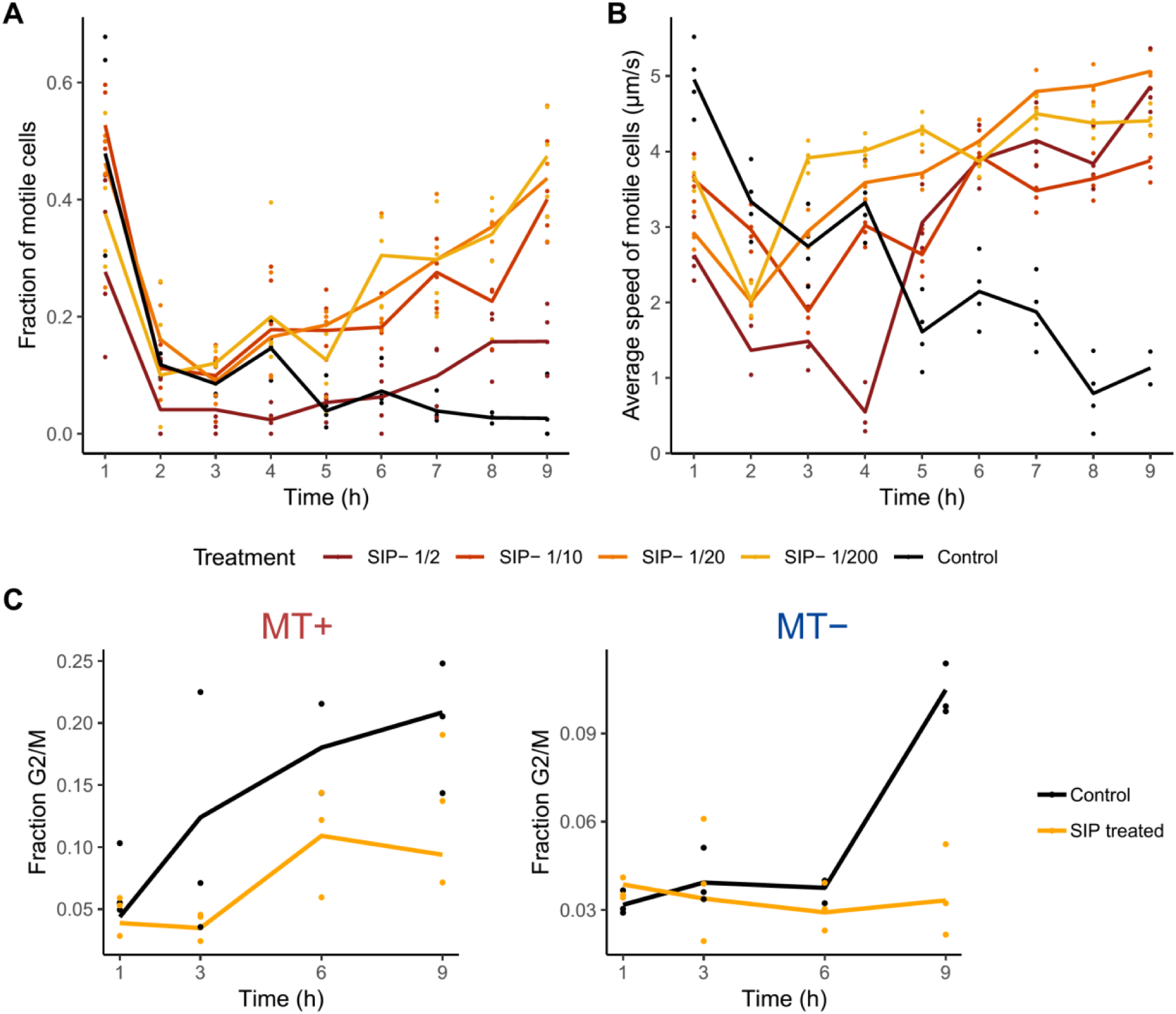
Physiological responses to SIP. **(A)** Fraction of motile MT+ cells over time for several dilutions of SIP- and a control treatment (n = 4). A significant increased motility is observed for 10x, 20x and 200x dilutions of SIP- at 7-9h (mixed model, p < 0.05). **(B)** Average speed of motile MT+ cells over time for several SIP- dilutions and a control treatment (n = 4). Significantly increased speeds are observed for all dilutions of SIP- at 7-9h. Note, that a high proportion of motile cells is observed after 1h in all samples, likely the result of a phototactic response following illumination after a prolonged period of darkness. **(C)** The fraction of cells in G2/M transition phase during the RNA-seq experiment for MT+, and for MT- (n = 3). For both MT+ and MT-, SIP induces a G1 phase cell cycle arrest, apparent by the significant lower fraction of cells in G2 and M phase after 3h and 9h (MT+) and 9h (MT-).

During S and G2 phase of the mitotic cell cycle, fission of both chloroplasts of *S. robusta* results in four daughter chloroplasts by the start of the M phase^46^. FtsZ (*Sro1409_g270150*) and dynamin-related protein 5B (DRP5B, *Sro814_g206290*) are key factors in the formation of a multi-ring structure that constricts the chloroplast during fission^47,48^. In control conditions, we observe an increased expression of FtsZ and DRP5B after 3h-6h of light exposure, coinciding with the timing of chloroplast division in *S. robusta*^*46*^. After exposure to SIP, expression of both genes was significantly repressed in both mating types (Figure 3), indicating that the cell cycle arrest results in an inhibition of chloroplast fission. Indeed, chloroplast division does not occur during sexual reproduction of *S. robusta*, as a result each gamete contains one chloroplast and the auxospore will inherit two chloroplasts^49^.

In the transcriptomic data, seven cyclins were downregulated in both mating types (Figure 3A). Six out of seven appear in a cluster of genes with peak expression late in the time series (6h, 9h and 10h, Supplementary Figure 7), suggesting that they are mitotic cyclins involved in G2/M transition^50^. To determine which cyclin family they represent, a maximum likelihood phylogenetic tree of *S. robusta* cyclins was constructed (Supplementary Figure 8). The repressed cyclins consist of 3 diatom-specific cyclins (dsCyc), 3 A/B type cyclins and 1 cyclin D (Supplementary Table 2).

Several genes involved in mitosis were downregulated in both mating types, including MAD2 (*Sro3109_g343920*), MAD3 (*Sro690_g187690*) and CDC20 (*Sro589_g171680*), which form the Mitotic Checkpoint Complex. Furthermore, we observe downregulation of 5 genes coding for subunits of the condensin complex, which play a central role in chromosome organization during mitosis and meiosis^51^ (Figure 3A).

Finally, at the end of each mitotic cell cycle, diatom cells produce a new valve in a silica deposition vesicle (SDV) prior to cytokinesis. As was previously shown^17^, treating cultures with SIP of the compatible mating type reduces the fraction of cytokinetic cells after 14h (Supplementary Figure 8, p < 0.0001 for both MT+ and MT-). Accordingly, we observed downregulation of genes known to be important for silica cell wall formation after treatment with SIP (Figure 3A). These include two silicic acid transporters: membrane proteins that are involved in the uptake of silicic acid from the environment^52^; two silicanins: proteins embedded in the SDV membrane^53^; as well as four frustulins: proteins which are found in the organic casing surrounding the cell wall^54^. Moreover, four genes making up various subunits of V-type ATPase complexes were significantly downregulated in both species upon treatment with SIP. These proton pumps play a role in biomineralization of silica by acidifying the SDV^55^.

While mitotic cell cycle genes were generally downregulated by SIP treatment, we observed an increase in meiotic gene expression. Surprisingly, one cyclin which is an SRB (*Sro299_g111470*, Figure 3) is upregulated rather than downregulated in response to the pheromone. Its expression pattern suggests it plays a role either in SIP induced physiological responses such as mate finding or in preparing the cell for meiotic cell cycle progression. The gene has presumably evolved independently from the sexually induced cyclins of other major eukaryotic clades, as phylogenetic analysis places the gene among diatom specific cyclins (dsCyc) (Supplementary Figure 8). The expansion of the cyclin family in diatoms compared to other members of the SAR clade^56^ might have been instrumental to allow the diversification of this sexually induced cyclin^57–60^.

To investigate the dynamics of other meiotic genes throughout the experiment, we explored the expression of a set of 42 known meiotic and bifunctional mitotic/meiotic diatom genes^5^, some of which were induced in response to sex pheromones in pennates^17,18^ and during sexual reproduction in the centric diatom *S. marinoi*^*42*^. Here, we found 10 meiotic markers significantly responding to SIP in at least one time point in both mating types (Figure 3). These include genes encoding DNA replication licensing factors MCM4, MCM6, MCM7, which are involved in the initiation of replication during the mitotic and meiotic S-phase^5^. Two homologues of SPO11 (SPO11-2 and SPO11-3) are upregulated in both mating types. SPO11 plays a role in the formation of double stranded DNA breaks during homologous recombination. In diatoms and plants, the SPO11-2 homologue was shown to be meiotic while SPO11-3 is involved in vegetative growth^5,18,61^. However, the observed upregulation of SPO11-3 in response to SIP in *S. robusta* and during sexual reproduction in the centric *S. marinoi*^*42*^ suggest that in addition to SPO11-2, also SPO11-3 might be involved in meiotic homologous recombination in diatoms. Three genes involved in DNA repair after the induction of double stranded breaks by SPO11 were upregulated in both mating types: MRE11, RAD50 and RAD51D^5^. Finally, we observed the upregulation of two genes not yet described in the meiotic toolkit of diatoms: Ataxia Telangiectasia Mutated (ATM, *Sro154_g070060*) which codes for a protein that controls double-strand break formation by SPO11 during meiosis^62^ and Replication Protein A (RPA, *Sro944_g222890*), involved in the binding of ssDNA during replication and homologous recombination^63^ (Figure 3). Interestingly, although meiotic genes are upregulated in response to SIP, the G1 phase arrest inhibits progression through meiotic cell cycle and we did not observe gametogenesis in MT+ cells attracted to diproline loaded beads. Accordingly, meiotic prophase markers and chloroplast rearrangements required for gametogenesis only take place after compatible cells form a mating pair^49^. We therefore hypothesize that a separate, local signal during cell pairing is required to break the G1-phase arrest and induce gametogenesis.

In response to SIP, 17 genes with a guanylate cyclase domain (GC) were significantly DE in both mating types, 8 of which form a bifunctional guanylate cyclase / phosphodiesterase (GC/PDE) fusion enzyme. Interestingly, the GC and PDE domains in these genes have contrasting functions, respectively synthesizing and breaking down the secondary metabolite cGMP. Although the genes with a GC domain do not show a general direction of regulation, all GC/PDE genes were upregulated (Supplementary Figure 10), including *Sro991_g228730*, which was previously described to be responding to SIP in *S. robusta*^*17*^. All significant GC/PDEs show the topology described by Moeys et al. (2016)^17^: a small N-terminal intracellular domain, an extracellular stretch and a long C-terminal intracellular part containing the GC and PDE domains. cGMP signaling appears to be a conserved response to sex pheromones in pennate diatoms, as guanylate cyclases are also upregulated in the pennate *P. multistriata*^*18*^.

Finally, three sex induced genes (SIGs), genes with unknown function induced during sexual reproduction in the centric diatom *S. marinoi* and the pennate diatom *P. multistriata*^*42*^, were upregulated in response to SIP (Supplementary Figure 11). SIG7 (*Sro587_g171310)* was significantly upregulated in both mating types. Protein domain analysis revealed the presence of a Homologous-pairing protein 2 (Hop2) domain in the *S. robusta* (IPR010776*), P. multistriata* (IPR010776) and *S. marinoi* (IPR040461) SIG7 sequence. Although Hop2’s role in homologous recombination would be in accordance with the observed expression, more work is needed to confirm SIG7 as a Hop2 homologue, as Hop2 is presumed to be absent in diatoms^5,42^. SIG9, meanwhile, is upregulated in response to SIP only after 10h of SIP treatment in MT- and SIG10a (*Sro637_g179400*) was significantly upregulated after 3h and 10h in MT-. Interestingly, protein domain predictions show that SIG10 consists of an unknown N-terminal domain, followed by one transmembrane helix and a C-terminal beta propellor domain in all three species (IPR013519, IPR011043 and IPR015943 in *S. robusta, P. multistriata* and *S. marinoi* respectively). This topology suggests a function as a receptor or in adhesion^64^. Since these three SIGs are upregulated in two pennate and one centric diatom species, they are interesting candidate marker genes for sexual reproduction in diatoms. Notably, we discovered 5 other SIG genes in the *S. robusta* genome which are not differentially expressed to SIP. If their function is conserved, we expect these genes to be upregulated during zygote or auxospore formation rather than during pheromone signaling, since the physiology of mate finding strongly differs between species.

### Responses to SIP specific for MT+

We identified 12 genes displaying a MT+ specific response to SIP (SRPs, Figure 4A-B). Seven of these genes lack known protein domains (Supplementary Table 1) and two others have limited functional annotation. The remaining SRP genes include a Myb-like/SANT-like transcription factor (*Sro94_g048900*), a metallopeptidase (*Sro489_g153380*), a gene encoding a transmembrane protein containing an EF-hand domain (*Sro1_g000260*) and a globin (*Sro2353_g324460*) (Figure 4C). The latter gene is especially interesting as, next to their oxygen binding capabilities, globins are increasingly implicated in redox signaling^65,66^. When exploring ROS signaling related genes, the conventional DE analysis uncovered an NADPH oxidase (NOX, Sro1572_*g283390*) with a pronounced MT+ specific response to SIP (Figure 3), catalyzing extracellular production of superoxide anions by moving an electron from NADPH through the plasma membrane to molecular oxygen. The observed NOX contains six transmembrane domains^67^ and an EF-hand domain suggesting a potential link with calcium signaling. As superoxide is cell impermeable, extracellular superoxide is most likely dismutated to H_2_O_2_, which can enter neighbouring cells for ROS signaling, e.g. through aquaporin channels^68,69^. NOX activity during sexual reproduction is a common theme in eukaryotes: it is required for gametogenesis and fertilization in plants^70^, it activates the mobility of spermatozoa in humans^71^ and is required for the formation of sexual fruiting bodies in the fungus *Aspergillus nidulans*^*32*^. Furthermore, gametophytes of the kelp *S. latissima* show female specific expression of NOX, suggesting mating type specific expression of NOX during sexual reproduction may be conserved among Stramenopiles^26^. Notably, enzymes from the NADPH producing PPP were upregulated in both mating types, potentially supplying NADPH to support NOX-mediated superoxide production.

The timing of the mating behaviour in *S. robusta* appears to be remarkably synchronized between mating types: we observed mate-searching behaviour of MT+ cells starting 6h-7h after treatment with SIP-, simultaneous with the first noticeable amount of diproline produced by MT-^15^ and the onset of responsiveness of MT+ to diproline (Supplementary Figure 9B). This behavioural change induced by SIP-was apparent from an increase in the fraction of moving MT+ cells (Omnibus test p < 0.001, Figure 5A) as well as from an increase in the speed of motile MT+ cells (Omnibus test p < 0.0001, Figure 5B). Thus, exposure to SIP does not only prime MT+ cells to become responsive to the attraction pheromone diproline, the increased motility of gametangia further increases the probability to encounter an immotile, diproline producing MT- cell. Moreover, pre-activation of cellular motility machinery by SIP- might explain the almost immediate attraction to a new diproline source^72^. As cell motility in raphid diatoms is achieved through the secretion of adhesive molecules that attach the cell to the substratum and provide traction for their gliding movement^73^, we performed BLAST searches to identify adhesive proteins containing a GDPH-domain, named after a conserved Gly-Asp-Pro-His amino acid motif^74^. Among 87 GDPH-containing proteins identified, four are upregulated in MT+ in response to SIP- treatment in the conventional DE analysis (Supplementary Figure 12). However, three out of four GDPH-domain containing genes are also significantly upregulated in immotile SIP+ treated MT- cells and one is upregulated uniquely in MT-. Therefore, as previously suggested, the GDPH domain containing proteins may play additional roles related to extracellular adhesion in addition to cell motility, such as mucilage pad, stalk and chain formation as well as cell-cell adhesion during mating^74^.

### Responses to SIP specific for MT-

A relatively high number (70) of SRMs, i.e. genes with a MT- specific response to SIP, was discovered (Figure 4A-B) of which 16 lack any functional annotation (Supplementary Table 1). Molecular functions of SRMs are diverse, including receptors, membrane channels, guanylate cyclases and other signaling enzymes. Among the SRMs are two E3 ubiquitin ligase genes (*Sro1305_g261220* and *Sro25_g017160*) representing the RBR and U-box family, respectively. Since ubiquitin ligases are important players in signaling by targeting downstream proteins for ubiquitination^75^, these genes are potential key regulators of MT- specific responses such as the production of diproline. Ubiquitin ligases also play a role in meiosis across eukaryotes by targeting proteins for proteasomal degradation^76^. However, a function in meiosis for the identified ubiquitin ligases is hard to reconcile with their mating type specific response, as meiosis occurs in both partners. Other SRMs include a gene with a DOMON domain which is a presumed haem or sugar sensor domain (*Sro7_g006210*)^77^ and a gene with homology to subunit D from the F0 complex of mitochondrial F-ATPase (*Sro89_g047110*) (Figure 4C). Interestingly, two SRM genes belong to an unknown and *S. robusta* specific gene family containing a zinc finger domain.

Our conventional DE analysis also shows that glutamate-to-proline conversion enzymes Δ1-pyrroline-5-carboxylate synthetase (*Sro2012_g310890*, P5CS), and Δ1-pyrroline-5-carboxylate reductase (*Sro216_g089310*, P5CR) were significantly upregulated after SIP treatment in MT- after 3h and 10h, while in MT+ both genes were not significantly responding (Figure 3B). Their MT- biased response supports the hypothesis that upregulation of P5CS and P5CR increases the cellular proline pool as a precursor for diproline biosynthesis^17^. Furthermore, a proline-tRNA ligase (*Sro27_g018090*) exhibits a strong significant upregulation uniquely in MT-, with expression levels exceeding 4000 CPM (Figure 3B). This enzyme attaches proline to transfer RNA (tRNA), which serves as a substrate for ribosomal protein synthesis, explaining its consistent expression in control conditions in this dataset (Figure 3B). The cyclodipeptide ring of diketopiperazines such as diproline is typically synthesized by either nonribosomal peptide synthetases (NRPS) or cyclodipeptide synthases (CDPS)^78^. Interestingly, CDPS require aminoacyl-tRNA as a substrate for the reaction instead of a free amino acid^79^. Thus, the observed MT- biased upregulation of a proline-tRNA ligase likely caters to the increased need for proline-tRNA^pro^ for CDPS dependent diproline biosynthesis. BLAST searches in the *S. robusta* genome revealed several candidate CDPS genes, and although some are expressed in all samples, none show upregulation uniquely in MT-. Thus, either an unidentified, transcriptionally controlled CDPS exists or one of the identified CDPS is non-transcriptionally regulated to be active only in SIP+ treated MT- cells with a cell size below the SST.

## Conclusions

This study provides a detailed overview of mating type specific expression in response to sex pheromones in pennate diatoms (Figure 6). To this end, we adopted a workflow that allows integration of independent RNA-seq datasets, prioritizing genes with shared and unique responses to SIP treatment. Gene expression changes closely correspond to physiological responses assessed by flow cytometry, mate searching behaviour motility measurements and chemo-attraction assays. We show that SIPs induce a G1 arrest which is reflected in a downregulation of essential genes involved in S-phase progression, chloroplast division, mitosis and cell wall formation. In contrast, multiple meiotic genes are upregulated in response to SIP, including the first sexually induced cyclin characterized in diatoms. Generally, our results highlight an important role for signaling through ubiquitination and cGMP, consistent with findings in other diatoms and brown algae. Additionally, combined evidence from the upregulation of a haem peroxidase, the NADPH producing pentose phosphate pathway, a globin and an NADPH oxidase (NOX) suggests that signaling through an oxidative burst is involved. Proline biosynthesis genes as well as a proline-tRNA ligase are upregulated uniquely in MT-, suggesting that the diproline synthesis pathway is cyclodipeptide synthase (CDPS) dependent. Furthermore, we report several unknown and often diatom- or species-specific SIP responsive genes. These genes should be prime targets for further research, as they likely encode novel proteins involved in the unique and largely unexplained sexual reproduction strategies of diatoms and *S. robusta* in particular. Unraveling their respective functions will lead to an improved understanding of the mechanisms underlying the fascinating life cycle of diatoms. Finally, our study identifies potential new marker genes which can be applied to natural situations to detect sexual events in diatom populations, providing a much-needed basis to document sexual reproduction in natural conditions.

**Figure 6:**
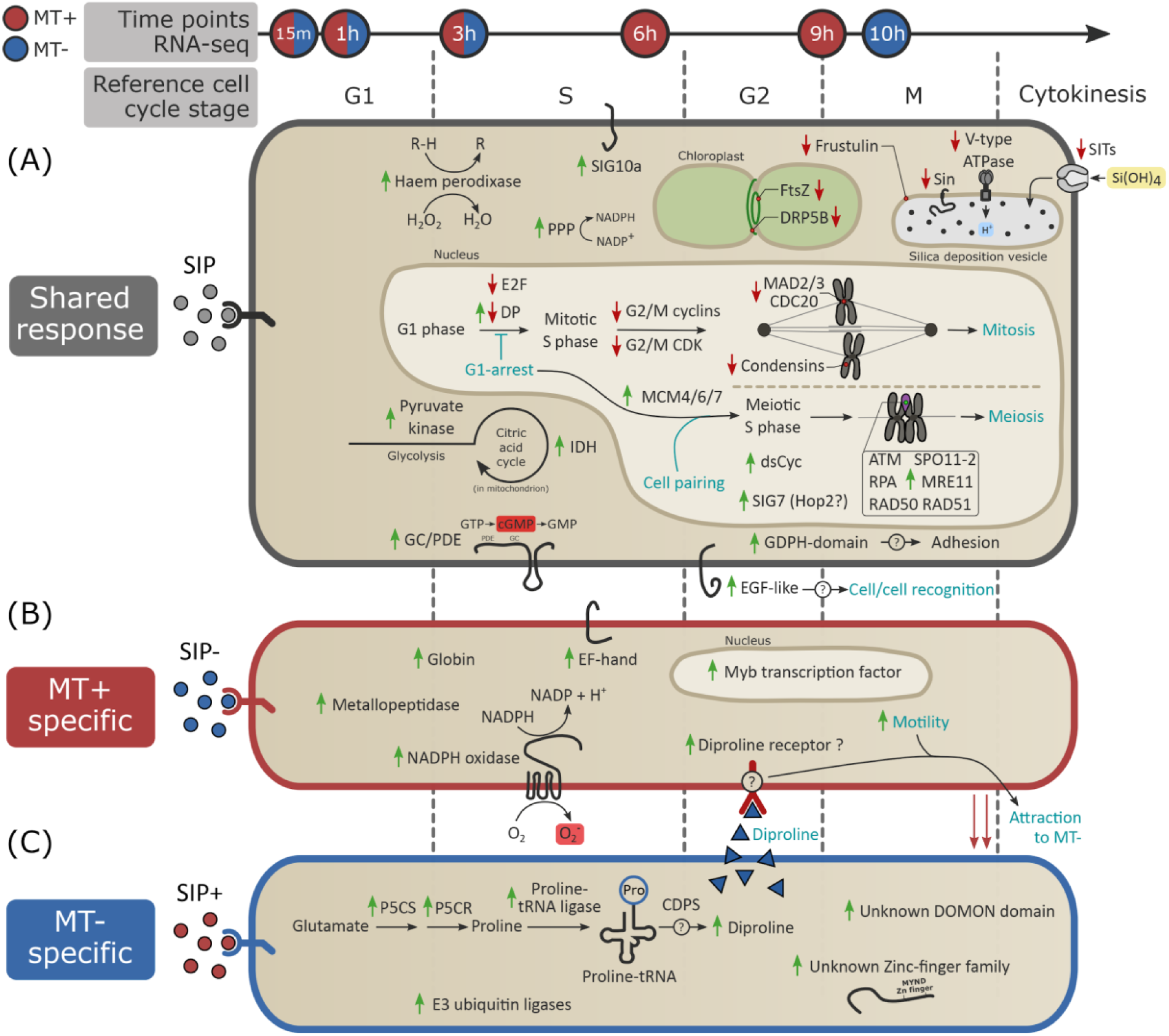
Overview of molecular and physiological processes in response to SIP in both mating types of *Seminavis robusta*. The arrow on top displays the harvest time of RNA-seq samples. Blue filled time points represent samples taken from MT-, red filled time points are samples from MT+ and red/blue filling indicates sampling for both mating types. The approximate timing of different cell cycle phases is shown below the arrow. **(A)** Cellular processes taking place in both mating types in response to SIP, **(B, C)** cellular responses to SIP unique for MT+ and MT-, respectively. Genes that are significantly up- or downregulated in response to SIP are depicted with a green or red arrow, respectively. Physiological events are indicated in cyan. PPP = pentose phosphate pathway, IDH = isocitrate dehydrogenase, P5CS = Δ1-pyrroline-5-carboxylate synthetase, P5CR = Δ1-pyrroline-5-carboxylate reductase and CDPS = cyclodipeptide synthase.

## Acknowledgements

We would like to thank Emmelien Vancaester for valuable contributions regarding gene family prediction. We are grateful to dr. Jeroen Gillard for providing some of the microscopic pictures of *S. robusta* sexual reproduction stages used in Figure 1. G.B. is supported by a Research Foundation Flanders (FWO) Aspirant grant (3F001916). K.V.d.B. is a postdoctoral fellow of the Belgian American Educational Foundation (BAEF), and is supported by the Research Foundation Flanders (FWO), grants 1S41818N and 1246220N. S.D.D. was supported by the Fund for Scientific Research – Flanders (FWO–Flanders, Belgium), grant No. G0D6114N and E.B. was supported by the research council of Ghent University (BOF/GOA No. 01G01715). N.P. was supported by the Deutsche Forschungsgemeinschaft (PO 2256/1-1). P.B. was supported by the Erwin Schrödinger fellowship from Austrian Science Fund (FWF) (J3692-B22) and FWO project G0D6114N. J.D. was supported by European Research Council grant ERC-ADG-670370. The research leading to the results presented in this publication was carried out with infrastructure funded by EMBRC Belgium – FWO project GOH3817N and funding from the research council of Ghent University (BOF/GOA No. 01G01715).

