## Supplementary figures and images for "Mating type specific transcriptomic response to sex inducing pheromone in the pennate diatom *Seminavis robusta*"

### Supplementary Figure 3_ SRBs.pdf

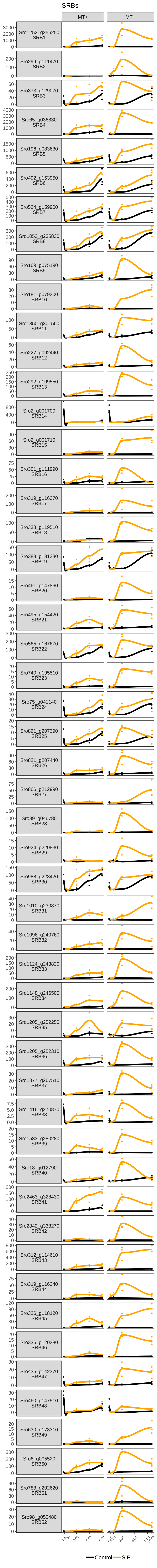

### Supplementary Figure 4_ SRPs.pdf

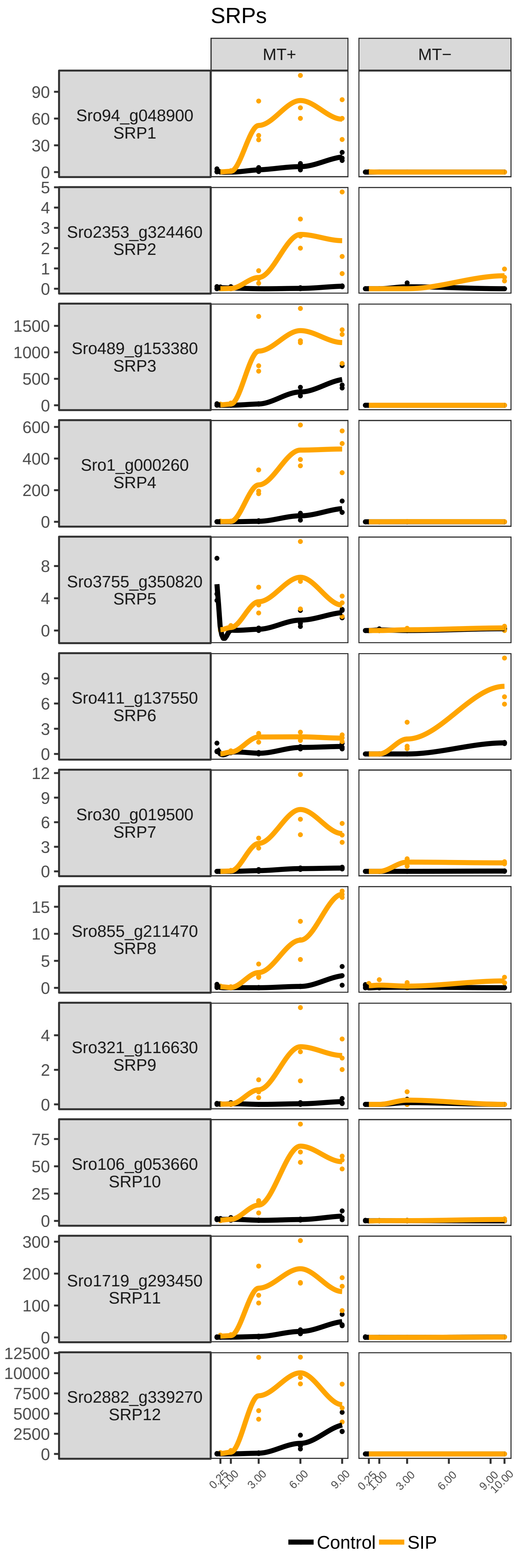

### Supplementary Figure 5_ SRMs.pdf

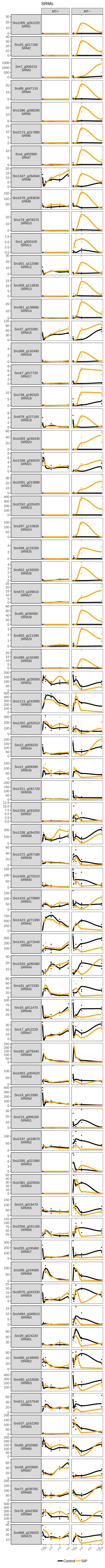
