## Supplementary Figures for "Mating type specific transcriptomic response to sex inducing pheromone in the pennate diatom *Seminavis robusta*"

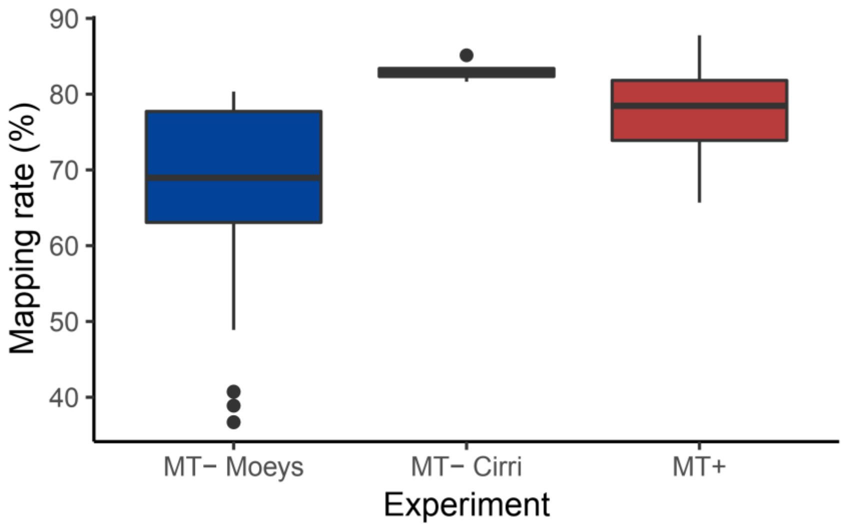


**Supplementary Figure 1:** Boxplots showing the distribution of the percentage of RNA-seq reads mapped with Salmon for each dataset. Data were retrieved from Moeys et al. (2016) (MT-)[1], Cirri et al. (2019) (MT-)[2] and new data presented here (MT+).

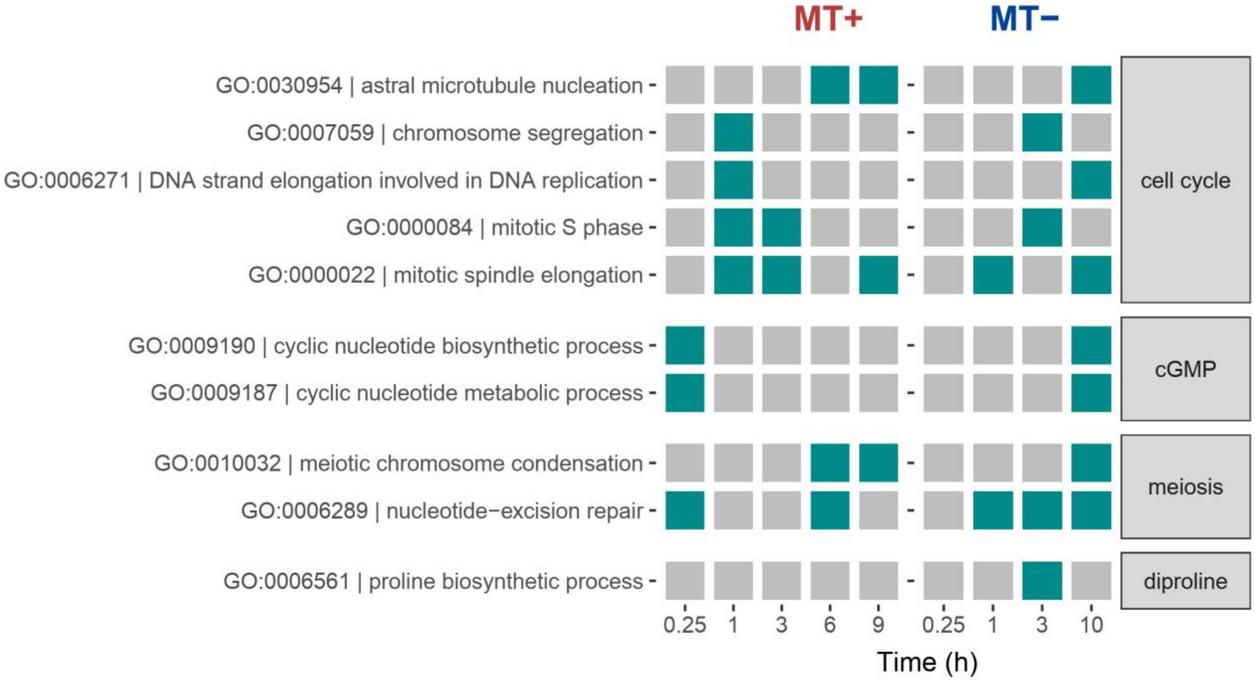


**Supplementary Figure 2:** Summary of gene ontology (GO) enrichment results using CAMERA[3] of selected known processes involved in mating. Gene ontology analysis was performed for each time point separately. Green boxes represent significant GO terms on a 5% FDR level for that specific time point. The left panel represents the results for MT+, while the right panel represents the results for MT-.

**Supplementary Figures 3, 4 and 5 are provided as separate pdf files in Supplementary.**

**Supplementary Figure 3:** Expression of all 52 key genes responding to SIP in both mating types (SRB genes), as identified by the integrative analysis. The x-axis represents time after SIP treatment. The left column represents gene expression in MT+ while the right column shows data from the two MT- experiments. The y-axis represents counts per million (CPM) for each gene, defined as the number of counts one would observe for a gene if each library were to be sequenced at a depth of one million.

**Supplementary Figure 4:** Expression of all 12 key genes responding to SIP only in mating type + (SRP genes), as identified by the integrative analysis. The x-axis represents time after SIP treatment. The left column represents gene expression in MT+ while the right column shows data from the two MT- experiments. The y-axis represents counts per million (CPM) for each gene, defined as the number of counts one would observe for a gene if each library were to be sequenced at a depth of one million.

**Supplementary Figure 5:** Expression of all 70 key genes responding to SIP only in mating type - (SRM genes), as identified by the integrative analysis. The x-axis represents time after SIP treatment. The left column represents gene expression in MT+ while the right column shows data from the two MT- experiments. The y-axis represents counts per million (CPM) for each gene, defined as the number of counts one would observe for a gene if each library were to be sequenced at a depth of one million.

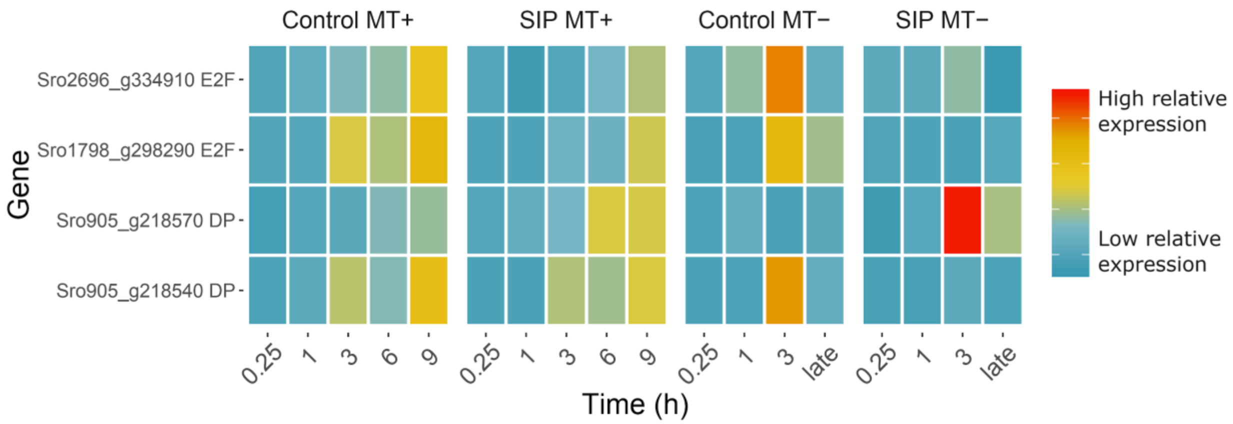


**Supplementary Figure 6:** Gene expression for two E2F and two Dimerization Partner (DP) transcription factors, which control S-phase progression during the cell cycle. Both E2F genes are downregulated upon SIP treatment, while one DP is upregulated (*Sro905_g218570*) and one is downregulated (*Sro905_g218540*). Genes are specified as row names and are coded according to their expression, amounting to counts per million (CPM) standardized to zero mean and unit variance for each gene in each mating type separately. Blue indicates low expression, while red indicates high expression.

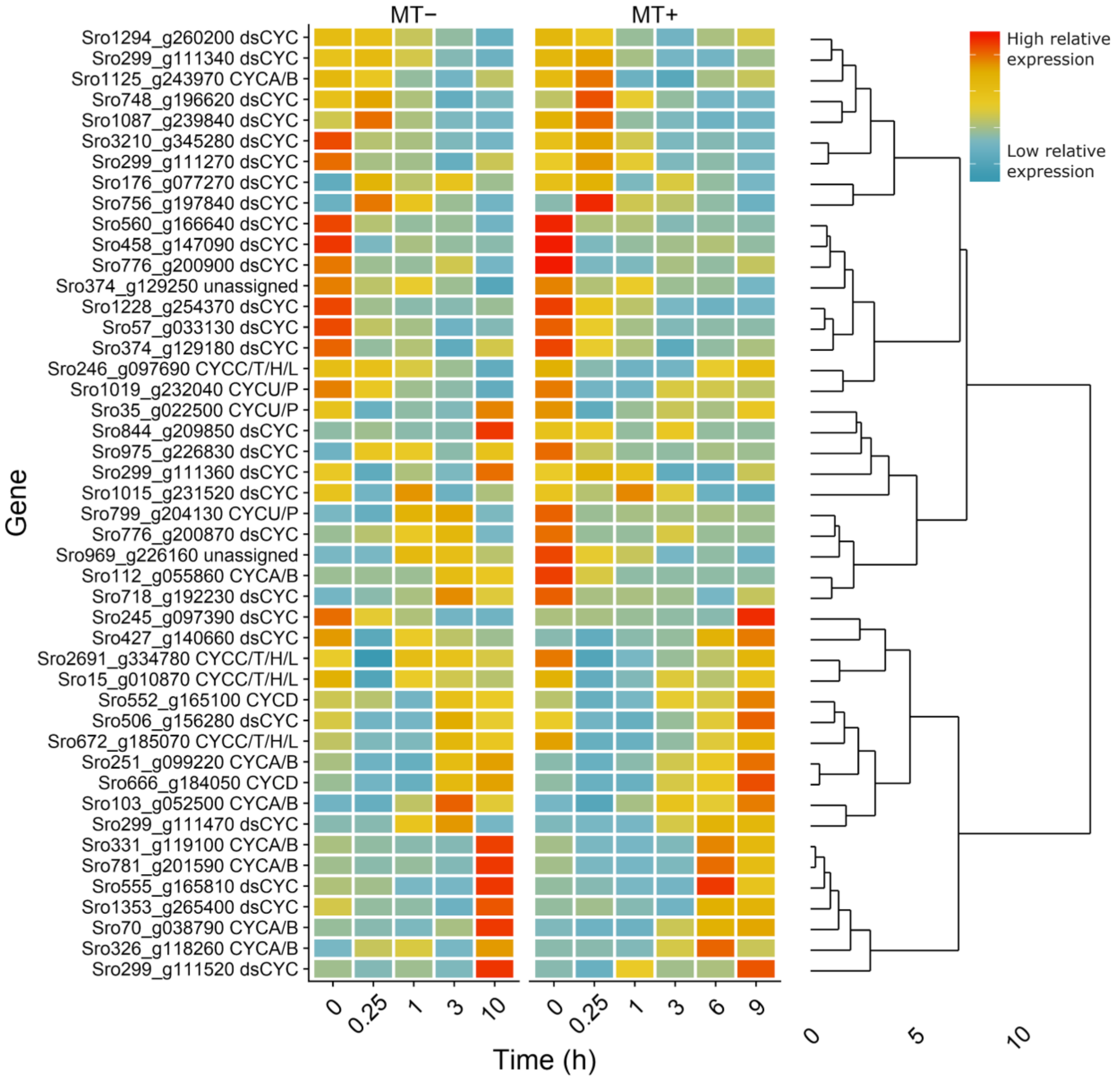


**Supplementary Figure 7:** Clustered heatmap of *Seminavis robusta* cyclins. Hierarchical clustered heatmap representing the expression of a selection of cyclins showing a change in expression during control conditions in datasets from MT- (left) and MT+ (right). Genes are specified as row names and are coded according to their expression, amounting to counts per million (CPM) standardized to zero mean and unit variance for each gene in each mating type separately. Blue indicates low expression, while red indicates high expression. The dendrogram structure of the hierarchical clustering of cyclin expression is plotted on the right side.

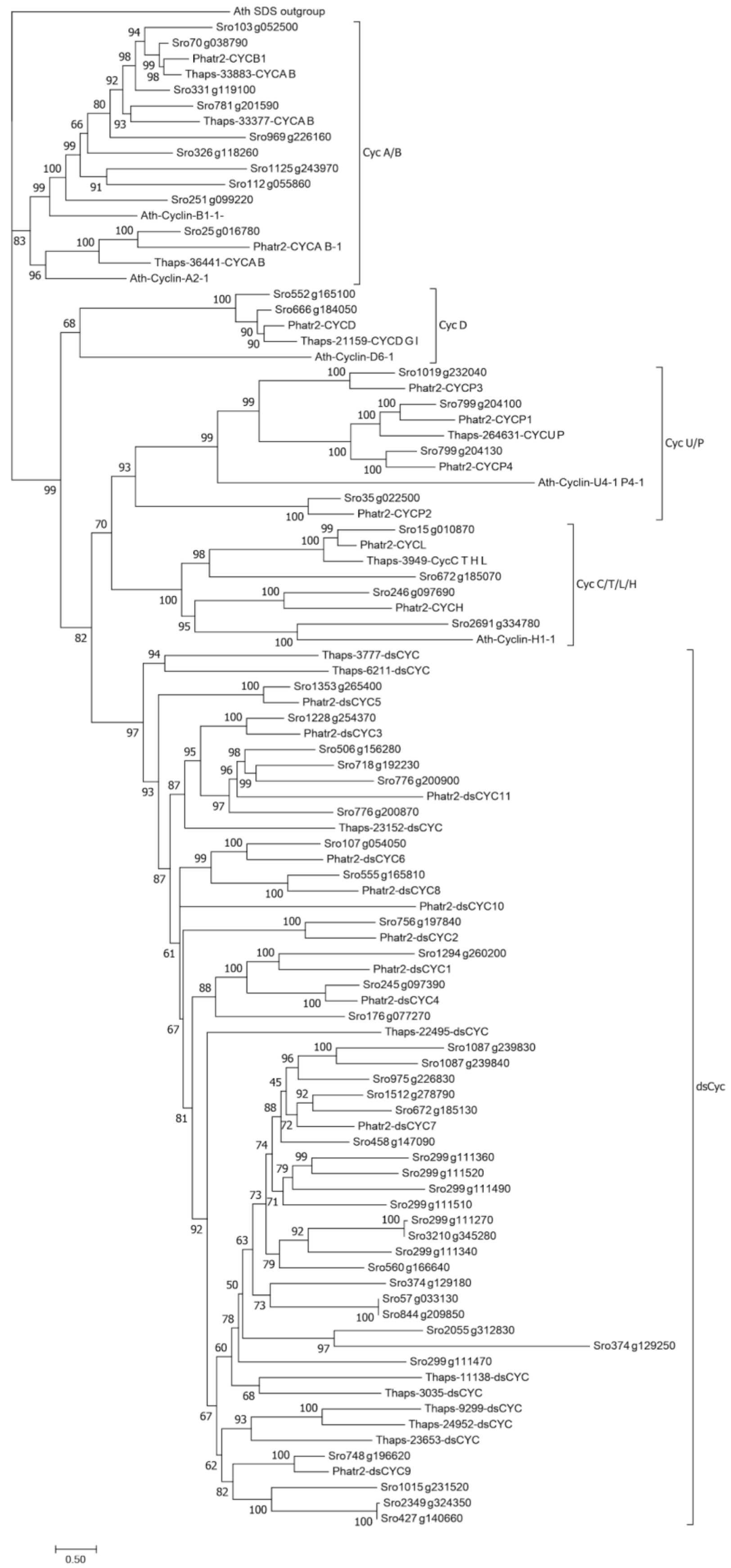


**Supplementary Figure 8:** Phylogenetic tree assigning *Seminavis robusta* cyclins to a cyclin family. Amino acid sequences of *S. robusta* cyclins were aligned with cyclins from model species *Phaeodactylum tricornutum*, *Thalassiosira pseudonana* and *Arabidopsis thaliana* using MAFFT v7.187. The phylogenetic tree was constructed using IQ-TREE. Numbers on the branches represent bootstrap support in percent, based on 1000 bootstraps. The outgroup is SDS (Solo Dancers) from *A. thaliana*.

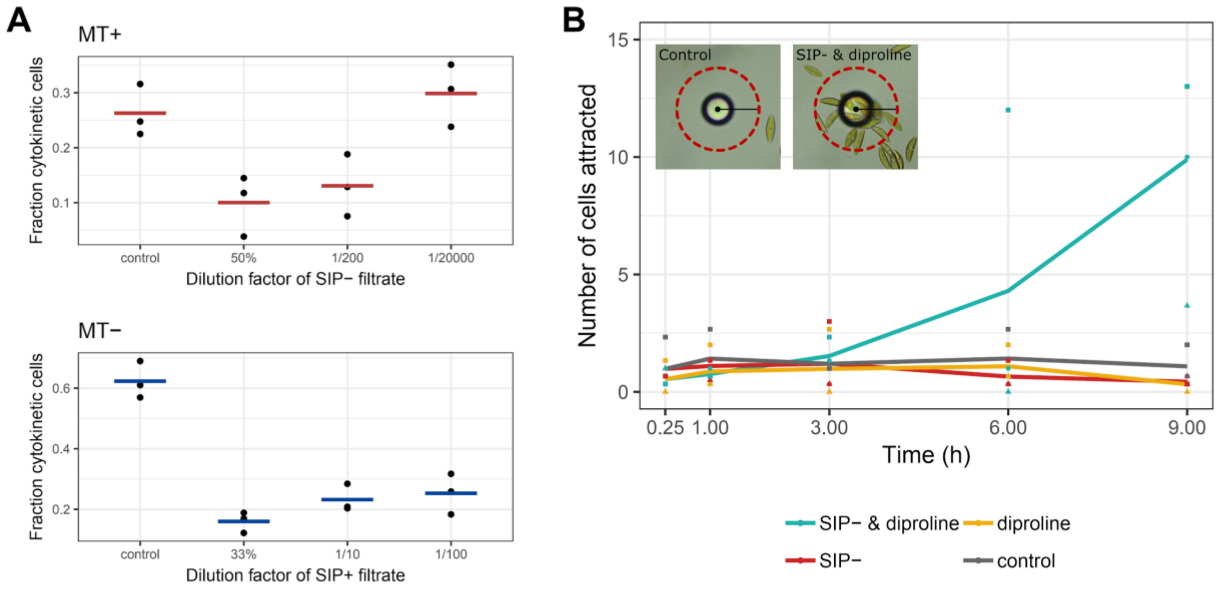


**Supplementary Figure 9:** Physiologic measurements testing the potency of SIP filtrate **(A)** The fraction of cytokinetic cells ("doublets") is given for cells of each mating type treated with different dilutions of SIP filtrate. The number of cytokinetic cells was determined 14h after first illumination and treatment with filtered pheromone. Points represent individual data points while horizontal lines show the mean for each treatment. **(B)** A bead attraction assay was performed to test the potency of SIP- filtrate used for the MT+ RNA-seq experiment. Beads were either uncoated or coated with a diproline-solution, while cultures were either untreated or treated with SIP- filtrate. The average number of cells attracted to each bead is given for different treatments. Time represents time since illumination and, if applicable, administration of SIP-. Points represent individual data points while lines show the mean for each treatment.

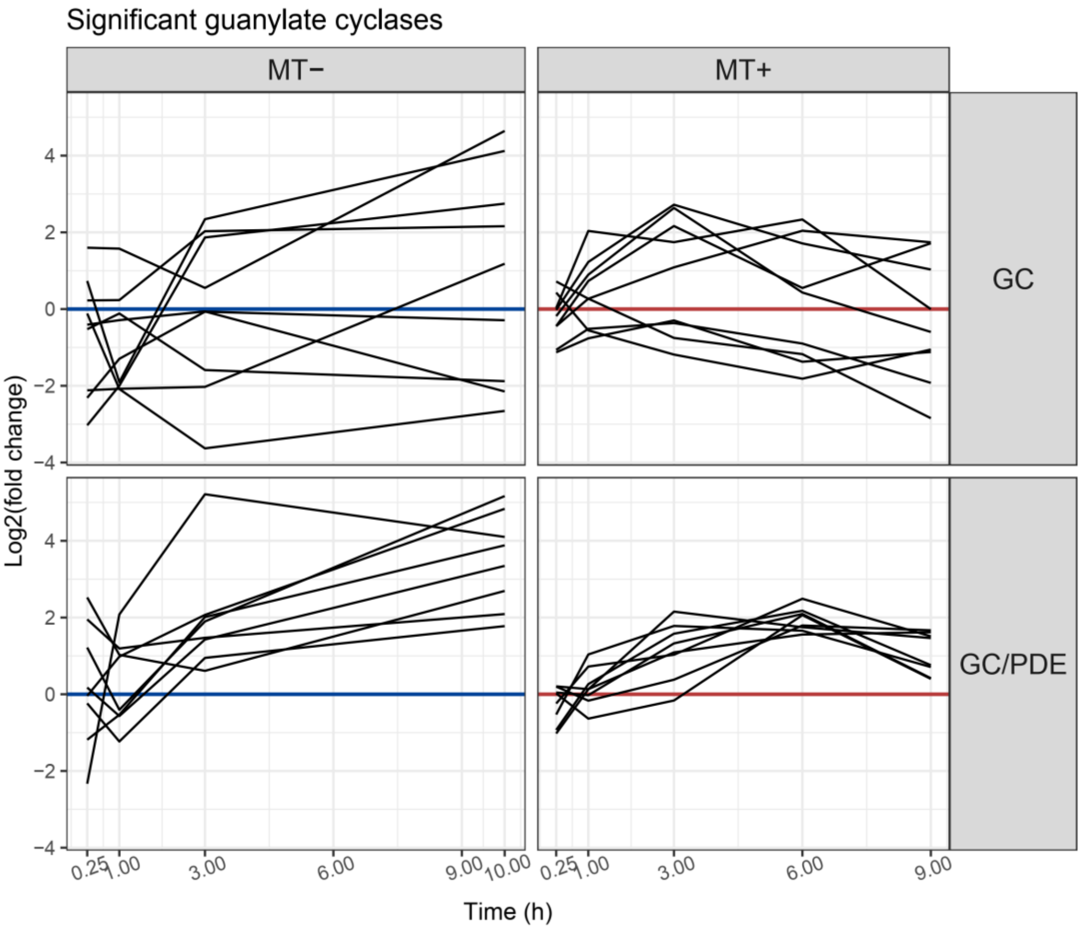


**Supplementary Figure 10:** Expression of GC and GC/PDE genes upon SIP treatment. Seventeen genes containing a guanylate cyclase (GC) domain are plotted. The x-axis represents time after SIP treatment and the y-axis shows the log2 fold change (SIP vs control) of average gene expression for each time point. The averages are connected for each gene using a straight line. The two columns represent the two mating types. In the top row, genes containing only a GC domain are plotted; with no obvious consistent fold change pattern. The bottom row shows genes containing both a GC and a phosphodiesterase (PDE) domain (GC/PDE bifunctional enzymes). Interestingly, all GC/PDE genes are upregulated upon SIP treatment.

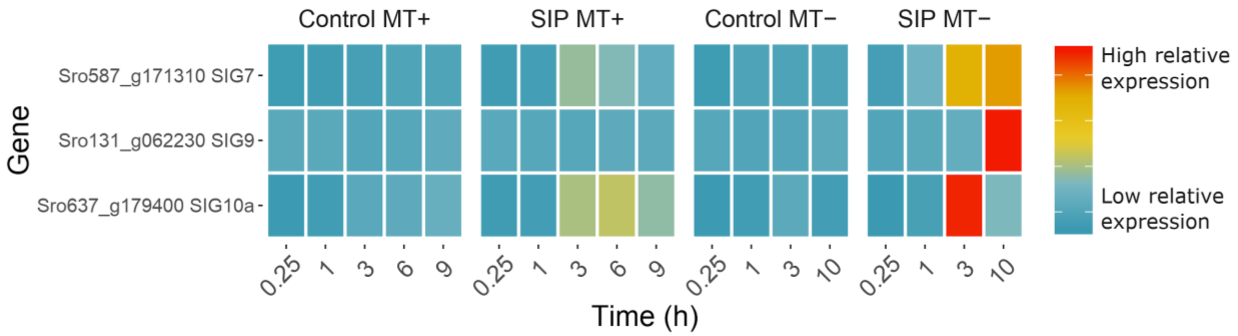


**Supplementary Figure 11:** Expression of three significantly differentially expressed Sex Inducing Genes (SIGs), as defined by Ferrante et al. (2019) [4]. SIG9 and SIG10a were only significantly upregulated in response to SIP in MT-, while SIG7 was significantly upregulated in both mating types. Genes are specified as row names and are coded according to their expression, amounting to counts per million (CPM) standardized to zero mean and unit variance for each gene in each mating type separately. Blue indicates low expression, while red indicates high expression.

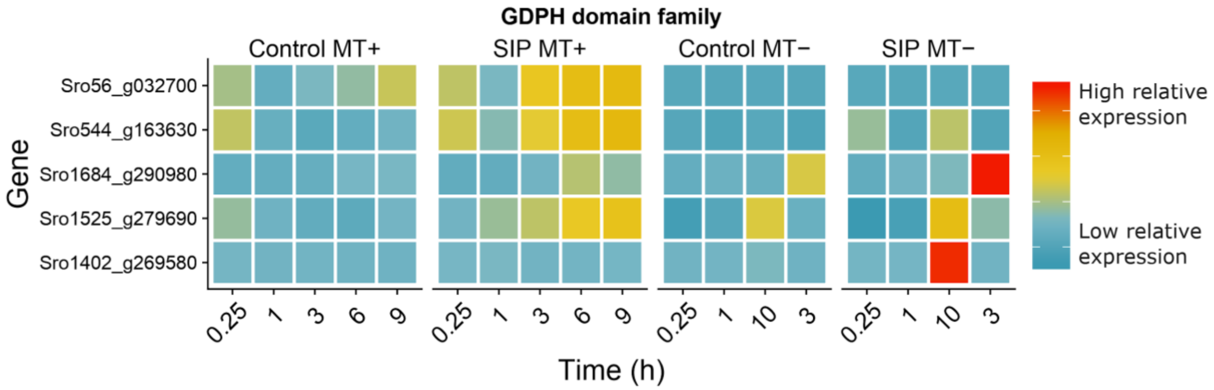


**Supplementary Figure 12:** Expression of five significantly differentially expressed genes containing a GDPH domain as described in Lachnit et al. (2019)[5]. The top gene is significantly upregulated only in MT+, the next three genes are upregulated in both mating types, while the bottom gene is uniquely upregulated in MT-. Genes are specified as row names and are coded according to their expression, amounting to counts per million (CPM) standardized to zero mean and unit variance for each gene in each mating type separately. Blue indicates low expression, while red indicates high expression.

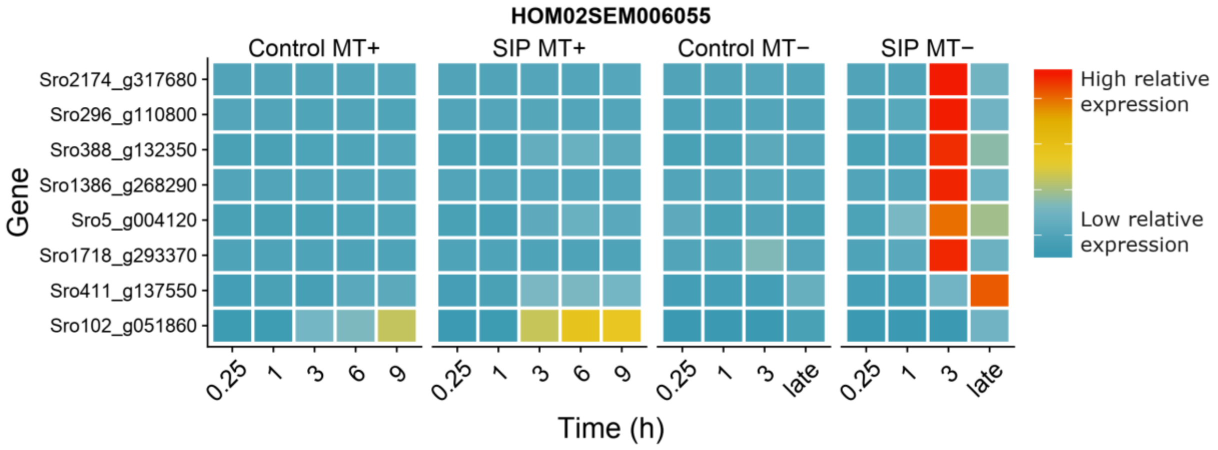


**Supplementary Figure 13:** Heatmap showing expression of significant genes from an unknown gene family of genes containing a zinc-finger domain. Eight genes of this family are plotted which are significantly differential expressed in at least one mating type. Two genes from this family were discovered as MT-specific genes (SRM) in our integrative analysis (*Sro2174_g317680* and *Sro1386_g268290*). In total, 7 genes are significantly upregulated only in MT- while the bottom gene is significantly upregulated only in MT+. Genes are specified as row names and are coded according to their expression, amounting to counts per million (CPM) standardized to zero mean and unit variance for each gene in each mating type separately. Blue indicates low expression, while red indicates high expression.

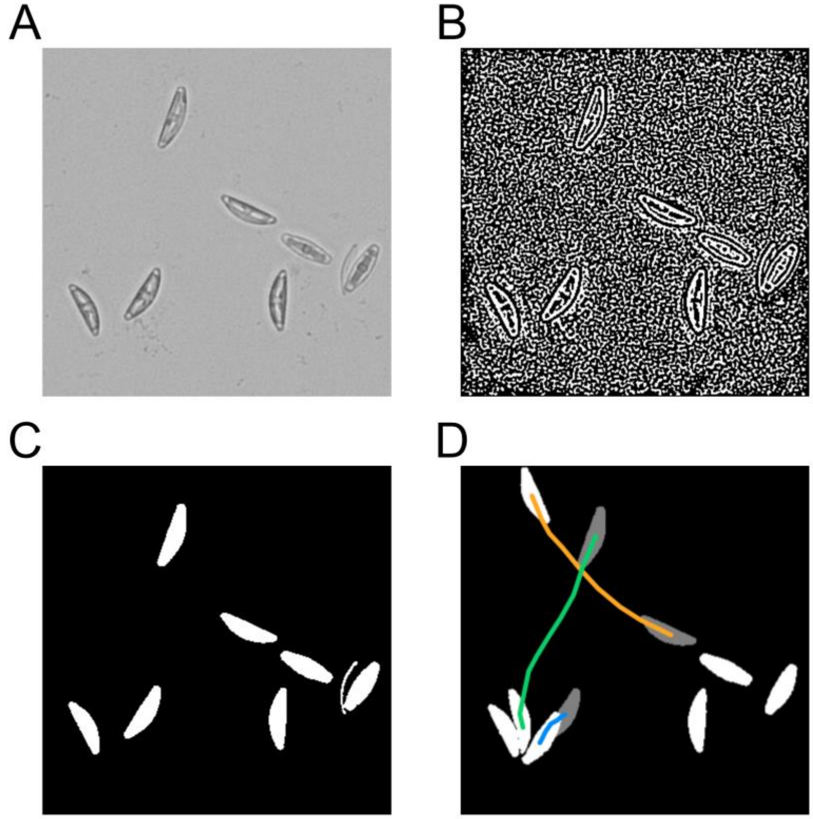


**Supplementary Figure 14**: Four different stages of image processing needed to track individual *S. robusta* cells. The images shown above are close-ups. **(A)** Movie frames were converted to grayscale images. **(B)** A LoG filter was applied to the grayscale images. The resulting image shows the diatom edges and background noise.**(C)** Black regions enclosed within a white diatom were filled. The properties of all resulting white regions were measured (e.g. area, solidity, minor axis length) and used to discriminate the diatoms from the noise. **(D)** For every frame, the centroids of the diatoms were calculated and connected to form a trajectory. Gray and white cells depict cell positions at frame 1 and 11 respectively. The colored lines depict trajectories of moving cells.

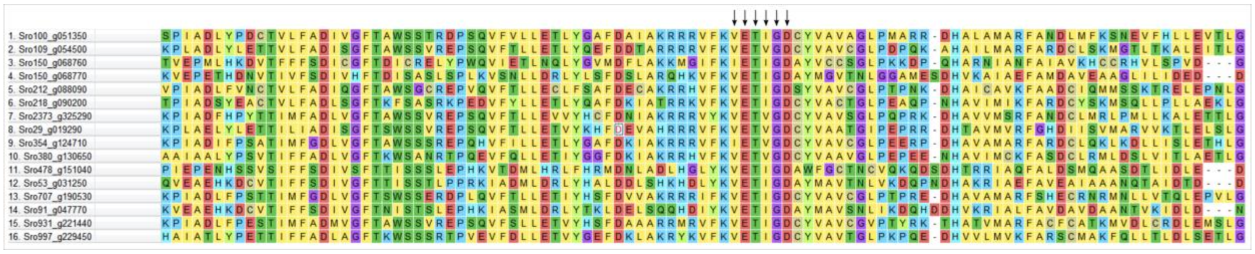


**Supplementary Figure 15:** Partial representation of an amino acid multiple sequence alignment performed with MUSCLE to confirm the substrate specificity of differentially expressed guanylate cyclases. Arrows represent a conserved guanine binding motif in the catalytic site of guanylate cyclases.
