## Supplementary Methods for "Mating type specific transcriptomic response to sex inducing pheromone in the pennate diatom *Seminavis robusta*"

#### Testing the potency of SIP- and SIP+ filtrate

The number of mitotically dividing cells in SIP treated versus control samples was counted to assess the potency of the filtrate. Late-exponential MT+ (85A) and MT- (PONTON34) cultures were synchronized by introducing a 36h dark period. Afterwards, respectively SIP- or SIP+ filtrate was added in three different concentrations (½, 1/200 and 1/20 000 for MT+ and ⅓, 1/10 and 1/100 for MT-). Control cultures received medium without SIP. Microscopic pictures were taken after 14h. We counted the proportion of cytokinetic cells using the Cell Counter plugin in FIJI (ImageJ). Statistical inference was carried out by modeling the expected fraction of cytokinetic cells using a quasibinomial generalized linear model (GLM) with logit link as a function of dilution. We first assessed the omnibus test, testing the null hypothesis that the expected fraction of cytokinetic cells is equal between each dilution and the control. Upon rejection of the omnibus test, a post-hoc analysis was carried out using the Multcomp package for R [1], performing Wald tests on a global 5% significance level to test the response of different dilutions vs. control within every time point.

A bead attraction assay was carried out to validate the attraction of MT+ to diproline after SIP-treatment. Oasis® solid phase Extraction cartridges (Waters Corporation) were equilibrated with 1 ml methanol and washed two times in MQ water, before loading 2 ml 0.00514 M synthetic diproline. Afterwards, the silica beads inside the cartridges were eluted in 2 ml MQ and subsequently diluted 1/50. Exponentially growing cultures of MT+ (strain 85A) were grown in a 24-well plate. After dark-synchronization of the cultures in G1-phase 50 µl of beads was added. Four different treatments were tested: (1) control, no conditioning with SIP-, no diproline on the beads; (2) conditioning with SIP-, no diproline; (3) no conditioning with SIP-, diproline on the beads; and (4) both conditioned with SIP and diproline loaded beads. For every well, a microscopic picture was taken from three random beads, 10 min after adding the beads. This procedure was repeated at 0.25h, 1h, 3h, 6h and 9h. Attraction was quantified by counting the number of cells in a 50 µm radius around the bead using image manipulation software ImageJ. For statistical analysis, we adopted a generalized linear mixed-effects model with Poisson distribution and log link function using the lme4 package for R (version 1.1-18-1 [2]). The number of cells was modeled as a function of treatment, time and their interaction effects as fixed effects. The effects of replicate and pseudoreplicate were added as random effects to account for multiple wells assessed for every replicate. Pairwise comparisons were carried out using Wald tests implemented in the glht function from the multcomp package for R. Attraction of beads was compared within every time point between each treatment and the control.

### Cell cycle analysis using flow cytometry

Cultures of MT+ (strain 85A) and MT- (strain PONTON34) were subjected to a 36h dark arrest to synchronize them in the G1 phase of the cell cycle [3]. Before re-illumination, half of the cultures were treated with a 1/10x dilution of SIP filtrate from the other mating type. Subsequently, 10 ml of culture was harvested and samples were centrifuged for 5 min at 1000 RPM. The supernatant was discarded and 10 ml ice cold 75% ethanol was added for fixation. The pellet was resuspended and stored in the dark at 4°C until further analysis. Later, fixed cultures were centrifuged for 5 min at 3000 RPM, after which the supernatant was replaced with 2 ml ice cold 75% ethanol. One ml of each sample was transferred to a 1.5 ml tube which was washed three times with PBS buffer. The fixed cells were treated with 1 µg/ml RNAse A for 20 min at 37°C, after which MT+ and MT- cultures were stained with propidium iodide (PI, 50µg/ml) and SYBR green (concentration 1x) respectively. Samples were filtered through a cell strainer with pore size of 70 µm and then analyzed on a Bio-Rad S3e cell sorter (BioRad laboratories, inc.). Data analysis was performed using the FlowCore 1.44.2 [4] and ggcyto 1.9.12 [5] packages for R. Debris was removed by gating and filtering on the forward scatter (FSC) versus side scatter (SSC) scatterplot, followed by a second gating step where unstained particles were removed. The number of cells in G1 and G2/M was identified by gating on the midpoint between both peaks on the histogram of intensities in the FL3 or FL1 channel respectively for MT+ and MT-. Statistical inference was carried out based on a quasibinomial generalized linear model (GLM) to account for overdispersion. The fraction of dividing cells was modelled as a function of treatment, time and their interaction effect. We accounted for the blocking structure in the experimental design of the MT+ experiment with a fixed effect for replicate. A global Wald test was performed as an omnibus test to assess any effect of the treatment in every time point. Upon rejection of the global test, a post-hoc analysis is carried out using Wald tests implemented in the multcomp R package [1] on a global 5% significance level, testing the response of treatment vs. control on the proportion of G2 cells within every time point.

### Assessing the effect of SIP- on MT+ motility

*S. robusta* MT+ (strain 85A) was grown at 18 °C under 20-25 µmol $m^{-2}s^{-1}$ fluorescent white light. Exponentially growing cells were inoculated in a 24-well multiwell plate (CELLSTAR®, Greiner Bio-one) filled with 1 ml of medium. The resulting cell density was 42.5 ± 16.6 cells $mm^{-2}$ (± SD) and cell length was 30.0 ± 1.8 µm (± SD, n = 30). At the end of the 36h dark-synchronisation period, 1 ml of SIP- filtrate was added in four different dilutions (1x, 5x, 10x and 100x) and a control without the pheromone was included by adding medium without SIP. Immediately afterwards, the cultures were transferred to the light. For each well, a 30-second microscopic movie was recorded every hour over the course of 9 hours. Quantification of motility over the 30 seconds period was performed with a custom MATLAB script (Supplementary Figure 14).

Every movie consisted of 11 frames and was exported as a multi-page tiff file. Every frame was imported in MATLAB as a gray-scale image and converted to a binary image. The centroids of the cells were used as cell coordinates and linked to a cell path by a MATLAB adaptation of John C. Crocker's particle tracking code [6]. Prior to the analysis, the dark corners of the image were corrected. For the analysis of the frames, the frames were filtered using a LoG-filter that acts as an edge detection filter. Removal of the generated noise and cell clusters yields a binary image consisting of single cells only. The output of the tracking code was a set of coordinates belonging to a path for every cell. The total length of the path was calculated and was used to determine if a cell was moving or not using a threshold-based procedure: since non-moving cells also generated a small path length, a set of movies without moving cells was generated by manually removing moving cells, and fed into the same algorithm described above. The resulting data was used to calculate a threshold for moving cells. The threshold was calculated to be the 95% quantile of the generated path lengths under no motility. Cells with a path length lower than the generated threshold were defined to be non-moving, those with a path length greater than the threshold were set to moving. For statistical inference we fitted a quasibinomial GLM to account for overdispersion. The expected fraction of motile cells was modeled as a function of treatment, time and their interaction effect. A global Wald test was performed to assess any effect of the treatment at every time point. Upon rejection of the global test, pairwise comparisons using Wald tests implemented in the glht function from the multcomp package for R[1]. The fraction of motile cells was compared within every time point between the 4 levels of dilution and the control.

### MT+ RNA-seq dataset experimental setup, RNA extraction, sequencing and processing

MT+ cells (strain 85A) were grown in 150 mL Cellstar® culture flasks (Greiner Bio-One GmbH, Austria). Two days before harvesting, the medium was replaced with fresh NSW + F/2 + antibiotics, and the density of the cultures was assessed using the F0 metric on a Maxi-PAM (Walz GmbH, Germany)[7] using parameters Intensity 7, Gain 3 and Damping 2. Cultures were subsequently diluted to an F0 value of 0.1. Before the start of the experiment, cultures were kept in the dark for 36h to ensure cell cycle synchronization in the G1 phase. Before re-illumination, half of the cultures cultures in 150 mL of medium were treated with 15 mL of SIP- filtrate. Fifteen minutes after addition of the filtrate, the light was turned on. At each of five time points (15min, 1h, 3h, 6h and 9h), two technical replicate flasks of SIP- treated and untreated cultures were harvested by filtration on a Versapor® filter with a pore size of 3 µm (Pall Corporation, NY, USA). The filters were subsequently rinsed with 1 mL PBS buffer, flash-frozen in liquid nitrogen and kept at -80°C until RNA extraction. The experiment was repeated on 3 different occasions to obtain 3 independent replicates.

Cells were scraped from the filters and lysed by adding 1 ml RLT lysis buffer containing 10µl $\beta$-mercaptoethanol. Silicon carbide beads (1 mm, Bio spec products Inc.) were added and the tubes were put in a beating mill (Retsch GmbH) for 30 minutes at 20 Hz. To extract RNA from the lysate, an RNeasy plant mini kit (Qiagen) was used according to the manufacturer's instructions, including an on-column DNAse treatment. One µl of each replicate was analyzed using a BioAnalyzer pico chip (Agilent technologies) to ensure RNA quality and concentration. The technical replicate of each repeat with best quality (BioAnalyzer RIN-value) and quantity (RNA concentration) was selected for sequencing. Library preparation and 2x75bp paired-end sequencing on the Illumina NextSeq500 platform occurred at VIB Nucleomics core (Leuven, Belgium http://www.nucleomics.be/). An average of 26.7±2.6 million reads was sequenced per sample. The samples were divided over two runs according to a randomized block design: all samples from the first repeat were assigned to the first run, and all samples of the second repeat were assigned to the second run, while samples of the third repeat were divided over the two runs in such a way that different treatments of one time point were within the same run. This block design allows an unbiased comparison of treatment effects, since all contrasts of interest can be assessed within a single run.

RNA-seq data generated in this study representing the response of MT+ to SIP- was complemented with existing data on the response of MT- to SIP+. We retrieved raw paired-end reads from Moeys et al. (2016) [8] for time points 15min, 1h and 3h, and from Cirri et al. (2019) [9] for a time point of 10h after re-illumination. In total, 54 samples were analyzed: 30 MT+ samples comprising 5 time points and 24 MT- samples comprising 4 time points. Quality control was performed on all samples using FastQC (Babraham Bioinformatics, under GPL3 license). Reads from Moeys et al. (2016) contained adaptor sequences which were trimmed with cutadapt v1.8 [10]. The sequences fragments were mapped using Salmon v0.9.1 (--gcBias for accounting for GC content bias) on the *S. robusta* gene models from Cirri et al. [9] (gene annotation v1.0, available at https://bioinformatics.psb.ugent.be/gdb/seminavis/Version1.0/). The average mapping rate was 77.7±5.7% for MT+, 66.9±13.9% for MT- from Moeys et al (2016) [8] and 83.1±1.2% for MT- generated by Cirri et al (2019) [9]. Note that the mapping rate is higher in the newer RNA-seq datasets as a result of improved data quality (Supplementary Figure 1).

Separate differential expression (DE) analyses were performed for each of the three datasets (i.e., the new MT+ RNA-seq time-series and two existing MT- RNA-seq datasets [8, 9]). Isoform-level abundances were imported in R using the tximport package (v1.8.0) [11] and aggregated to the gene level. Independent filtering [12] was performed by retaining genes with at least 1 count per million (CPM) in at least three samples. Differences in sequencing depth and RNA population were corrected for by adopting TMM normalization [12, 13] and including the natural logarithm of effective library sizes as offsets to the model. Negative binomial GLMs were estimated for every gene with edgeR [14], and gene expression was modelled as a function of an interaction between treatment and time. For the novel MT+ data and the MT- data from Moeys et al (2016) we account for technical effects by incorporating main effects of run and replicate, to respectively account for the sequencing run effects and block design imposed by the data generation structure (i.e., the data for different replicates may be gathered in different weeks). DE between treatment and control at every time point was tested for using likelihood ratio tests (LRT). Since interpretation and biological validation occur at the gene level, we adopted stage-wise testing to control the gene-level false discovery rate (FDR) [15]. To investigate transcriptional trends at each time point, a GO enrichment analysis was carried out (Supplementary Table 3).

### Functional annotation of *Seminavis robusta* genes

Functional annotation for all *S. robusta* genes was derived using three different approaches: (i) InterProScan v5.3 [16] was executed to search for matches against the InterPro protein signature databases; (ii) AnnoMine [17] was applied for consensus gene functional annotation retrieval from protein similarity searches (using DIAMOND v0.9.9.110 [18], maximum e-value 10e-05) against the Swiss-Prot database [19]; (iii) eggNOG-mapper [20] was used in DIAMOND mapping mode, based on eggNOG 4.5 orthology data [21]. To compute gene families, an all-against-all protein similarity search was performed using DIAMOND (maximum e-value 10e-05, max 4000 hits), after which the protein sequences were clustered into families using TRIBE-MCL v10-201 [22]. The set of species used for this analysis is shown in Supplementary Table 4.

The presence of specific conserved motifs was confirmed using multiple sequence alignment in CLC Main Workbench (Qiagen) for GDPH-domain containing proteins [23], frustulins [24], sin1-homologues [25], silicic acid transporters and guanylate cyclases (GC) [26]. To verify the specificity for guanine over adenine, the amino acid sequence of the catalytic domain of GCs was verified using multiple sequence alignment with MEGA7 [26] (Supplementary Figure 15) and the presence of transmembrane domains was determined using Phobius [27]. To characterize the cyclin family in S. robusta, a phylogenetic analysis was performed using 56 putative S. robusta cyclins and a selection of cyclins from P. tricornutum, T. pseudonana and A. thaliana. The A. thaliana SDS (At1g14750) amino acid sequence was used as an outgroup. Multiple sequence alignment was carried out with MAFFT v7.187 [28] after which poorly aligned positions were trimmed using Trimal V1.4.1 (-gt 0.1) [29]. The phylogenetic tree was constructed using IQ-TREE v1.7 (-bb 1000 -mset JTT,LG,WAG,Blosum62,VT,Dayhoff -mfreq F -mrate R) [30].

To identify SIGs in the genome of *Seminavis robusta*, BLASTp searches were carried out based on amino acid sequences from *Skeletonema marinoi* and *Pseudo-nitzschia multistriata*. Afterwards, amino acid sequences from each species were used for InterPro domain prediction [31].

### Gene set enrichment analysis

Gene ontology (GO) enrichment analyses were adopted to interpret transcriptional changes observed at every time point for each mating type. Competitive gene set testing was carried out on GO terms predicted by the EggNOG-mapper with CAMERA [32]. Briefly, CAMERA tests whether a set of genes is highly ranked for the hypothesis of interest as compared to other genes outside the gene set, while accounting for inter-gene correlation. Using CAMERA, a set of GO terms was defined for every time point of every mating type that are enriched in significant genes at the 5% FDR level. Enriched GO terms were summarized using REVIGO [33] using a similarity cut-off value of 0.5 and the SimRel score as a similarity measure[34]. REVIGO-summarized terms involved in processes of interest were selected and their enrichment was verified in other time points from both mating types. We selected GO terms from processes of interest and results were plotted in a heatmap showing the enrichment of each term in each time point and mating type combination (Supplementary Figure 2).

### Integrative analysis

SRBs, i.e. genes with a response to SIP in both mating types, were selected by testing whether their fold change was significantly higher than 3 or lower than 1⁄3 using a test for DE relative to a fold change threshold [8, 35]. To control the gene-level false discovery rate (FDR) on a 5% level for each dataset, a stage-wise testing approach was used [15] based on Sidak p-value aggregation. We first select these genes in the MT- dataset, and subsequently analyze this subset in the MT+ dataset using the same approach, i.e., we test against the same fold change threshold in at least one time point on a 5% gene-level FDR for the subset of MT- significant genes. This workflow guarantees FDR control on the full set of SRB genes.

A two-step procedure was used to discover genes that respond to the pheromone in only one mating type (SRMs and SRPs). In a first step non-responsive genes were detected for each mating type through equivalence testing [36] by selecting genes whose fold change was significantly contained within an equivalence interval ranging from ⅓ to 3 on a 5% FDR level. The equivalence tests were performed using the two one-sided tests (TOST) procedure for every contrast. To test for genes that are equivalent across all 3 time points, we controlled the FDR on the maximum p-value across all time points for a given gene. Since genes that are not expressed in a particular mating type (MT) can also be considered to be non-responsive, equivalent genes were subsequently merged with lowly expressed genes (CPM < 1 in > 3 samples) that were previously filtered out of the analysis. Next, we set out to identify which of the non-responsive genes in one MT were responsive in the compatible mating type. We identified genes with a fold change significantly higher than 3 or lower than ⅓ in at least one time point using an omnibus test on a 5% FDR level, producing lists of genes that are responsive in one MT, but non-responsive in the other. Expression in control and SIP treated conditions was plotted for a selection of candidate genes from each class (Figure 4).

### Visualisation of genes of interest

We investigated the response over the entire time interval of genes of interest using heatmaps which show the expression of significant genes in each time point for each mating type. Average counts per million (CPM) were calculated for every treatment*time combination. To improve visualization, average CPM were scaled separately for each mating type to zero mean and unit variance within each gene, except for supplementary heatmaps showing a mating type specific response, where scaling was performed over both mating types for each gene. All heatmaps were plotted with the ggplot2 package [37] using a gradient palette from the R package ‘wesanderson' [38]. To explore the synchronized expression of cyclins in the control samples during the time series, cyclins with changing expression in both mating types over time were selected (maximum CPM must exceed 2 times the minimum CPM), ordered based on their expression profile using hierarchical clustering in R.
